# Apolipoprotein-E transforms intracellular Amyloid-β oligomers to a more toxic state

**DOI:** 10.1101/2023.09.03.556061

**Authors:** Arpan Dey, Aditi Verma, Uchit Bhaskar, Bidyut Sarkar, Mamata Kallianpur, Vicky Visvakarma, Anand Kant Das, Kanchan Garai, Odity Mukherjee, Kunihiko Ishii, Tahei Tahara, Sudipta Maiti

## Abstract

It is poorly understood why ApoE variants are major genetic risk factors in Alzheimer’s disease (AD), which is associated with the aggregation of amyloid beta (Aβ). Here we directly image specific changes in small Aβ oligomers in rat brain cells that correlate with the cellular ApoE content. An inhibitor of Aβ-ApoE interaction suppresses this change and concomitantly reduces Aβ toxicity in a dose-dependent manner. Single-molecule techniques show changes both in the conformation and the stoichiometry of the oligomers. hiPSC-derived neural stem cells from Alzheimer’s patients also show similar changes. Interaction with ApoE therefore changes the oligomeric state, membrane affinity, and toxicity of Aβ oligomers, and can be directly read out in live cells. Our findings suggest a rapid and quantitative assay for AD drug discovery.

**One-sentence summary:** ApoE causes specific toxicogenic modifications of Aβ oligomers, and these changes can be directly imaged in live cells.

## Main Text

Alzheimer’s disease (AD) is associated with the aggregation of amyloid-beta (Aβ) and phosphorylated tau (p-tau) proteins in the brain(1). It presents a significant health burden, but limited understanding of the toxic pathway has hampered the search for a cure. Apolipoprotein E (ApoE), particularly the ApoE4 allele, is the strongest non-Aβ risk factor associated with late-onset of AD(2). Aβ and p-tau aggregates are directly observable in AD patients’ brains, but their interaction with ApoE (if any) remains less well established(3–6).

A plausible interaction of ApoE with Aβ has been a focus of multiple solution-state and intracellular studies(7). However, these have produced inconsistent results(8,9), possibly due to variations in the lipidation state of ApoE(10) and the oligomeric state of Aβ between different experiments. Strong evidence for a direct interaction of ApoE with Aβ comes from peptides that mimic stretches of Aβ and can block such interactions *in vitro*, and can also significantly reduce Aβ toxicity in cells(11,12). However, there is no direct read-out of ApoE-Aβ interaction in living cells. ApoE need not interact directly with Aβ to induce toxicity (13,14). Indeed, ApoE has toxic interactions with gamma secretase(15) (an enzyme that is in the pathway of Aβ production) and also with tau(3,16).

An understanding of ApoE-Aβ interaction, if any, can provide an essential foundation for exploring the toxic pathway in AD and offer a route to new therapeutic approaches. However, the paucity of direct, quantitative readouts of ApoE-Aβ interactions in living cells hampers such understanding. To address this challenge, here we probe fluorescence lifetime (called”lifetime” henceforth), which is one of the most sensitive reporters of local molecular interactions that allow quantitative spatio-temporally resolved measurements in live cells. Even if ApoE-Aβ interaction occurs only with a specific subset of Aβ aggregates(17) and/or if it is transient, fluorescence lifetime imaging microscopy (FLIM) is likely to report it. Here we find that FLIM data bears a quantifiable signature of intracellular ApoE-Aβ interaction, which we further validate by single-molecule experiments on cell extracts. This signature also strongly correlates with Aβ toxicity.

### Fluorescence Lifetime Imaging reveals a signature of intracellular interaction between ApoE and Aβ

We performed FLIM of an N-terminal rhodamine-B tagged Aβ (**RAβ**) in different cell types containing different natural abundance of ApoE, and correlated the change of fluorescence lifetime to the ApoE concentration present in each. Several different cell types were chosen to span a range of intracellular ApoE content(18,19). These included primary cortical neurons and astrocytes obtained from the brains of WT Sprague Dawley pups P0-P4 (details in **SI 1, figure S1** and identification of astrocyte in **SI 2, figure S2**), RN46A (a rat neuronal cell line), and HeLa cells (of human somatic origin). Cells were incubated with freshly prepared 100 nM RAβ which was taken up by the cells (20) and the lifetime of the N-terminus rhodamine label was monitored via FLIM (schematic in **Fig 1A**, and **SI 3**). At this concentration range, small oligomers (mostly <10-mers) are spontaneously formed, but no further aggregation occurs (21). These oligomers have been characterized earlier by us in terms of size, stoichiometry and residue level conformations(21–26).

**Figure 1.**
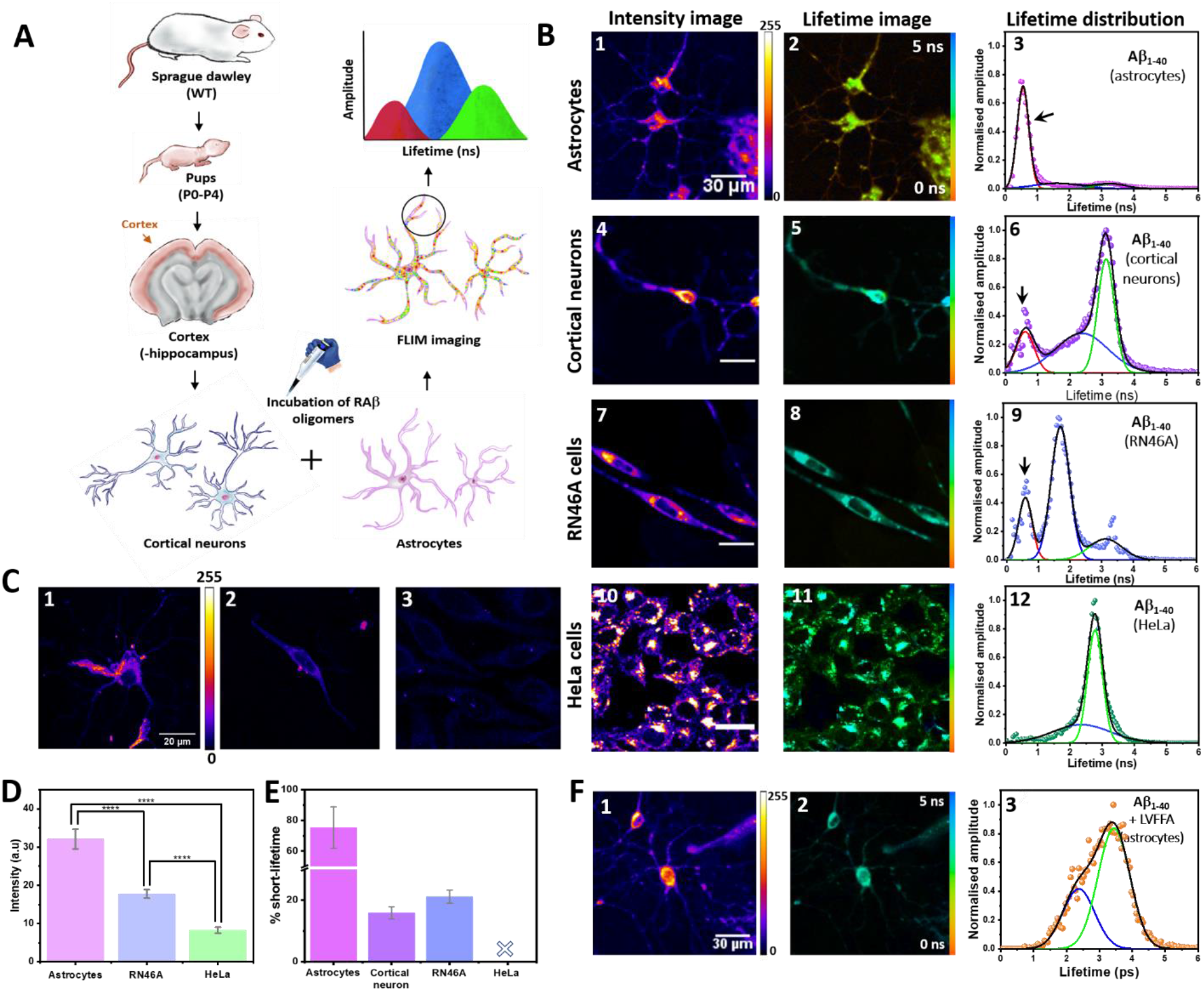
Fluorescence lifetime of Aβ N-terminal label correlates with intracellular ApoE-Aβ interaction: **A)** Schematic representation of the experiment. **B)** Fluorescence Intensity, lifetime, and average lifetime distribution of RΑβ oligomers in astrocytes (1,2, and 3), cortical neurons (4,5, and 6), RN46A cells (7,8,9), and HeLa cells (10, 11, and 12). The average lifetime distribution is fitted using three Gaussians (description in **SI 5, figure S3**), representing short, intermediate and long lifetimes, with their peaks allowed to vary within a certain range. The first component has a short-lifetime (red, peak at 530 ps ± 20%), second component has a lifetime similar to RΑβ in buffer (or to free rhodamine B for RN46A cells) (blue, peak at 1974 ps ± 20%), and the third component has a longer lifetime (green, peak at 3270 ps ± 20%) similar to RΑβ in lipid membranes (data in **SI 6, figure S5**). Lifetime distribution shows the short-lifetime population gradually decreases from astrocytes to RN46A and is absent in HeLa. Lifetime distribution data represents cumulative average distribution from three independent measurements of all the cell lines. Each measurement represents three dishes (six different fields per dish) from each cell type. **C)** Quantification of ApoE in different cells using ICC. Primary antibody – recombinant anti-ApoE antibody /abcam EPR19239 - reacts with mouse, rat, and human ApoE, (specificity against rat ApoE checked by Western Blotting in SI), secondary antibody Goat Anti-Rabbit IgG H&L (Alexa Fluor(R)647) pre-adsorbed. (1) ICC Intensity images of astrocyte, (2) RN46A cells, and (3) HeLa cells. **D** Average intensity of secondary antibody per unit area from the Astrocytes, RN46A, and HeLa cells (data averaged over three replicates each having three slides per replicate. Number of cells analysed are 32 astrocytes, 58 RN46A, and 61 HeLa cells). Controls (ICC without primary antibody shows no non-specific binding of secondary antibody, **in SI 8, figure S7**). Quantification shows maximum expression of ApoE in astrocytes and least in HeLa. **E)** The percentage of short-lifetime in different cells having different ApoE content (area of the short-lifetime Gaussian normalised by the total area under the curve). A qualitative similarity between cellular ApoE content and short-lifetime indicates an ApoE mediated effect on the lifetime of N-terminus rhodamine B label of Αβ. **F)** Fluorescence Intensity (1), lifetime (2), and average lifetime distribution (3) of RΑβ oligomers in astrocytes in the presence of 30 µM of LVFFA (inhibitor of Αβ−ΑpoE interaction?. The absence of the short-lifetime in astrocytes, and the emergence of the medium and the long components show that interaction between ApoE and RΑβ is required for generating it (intensity and lifetime calibration bar and scale bar for all the images in **B** are same (8 bit images, lifetime distribution from 0-5 ns, and 30 µm) and in **C**, the calibration bar is same, but the scale bar for all the images is 20 µm. Error bars in figures **D** are mean ± SEM, significance of difference is performed by T-test, P < 0.0001); error bars in figure E are errors of the fit.

Fig **1B** shows the fluorescence intensity and lifetime images in different cells and the corresponding average lifetime distributions. The lifetimes were analysed using a quasi-continuous distribution (see **SI 5, figure S3, table S5-9**) which showed a considerable width reflecting heterogeneity in the samples. However, independent fits of individual datasets, followed by global Gaussian fits of the average lifetime distributions, yielded three major populations with average lifetimes in the range of 0.53 ± 0.10 ns, 1.97 ± 0.39 ns, and 3.27 ± 0.65 ns respectively (fit parameters, independent fits, and choice of lifetime range described in **SI 5, table S5**). The shortest lifetime component was absent in HeLa cells. These three lifetime components are subsequently called the ‘short’, ‘intermediate’, and ‘long’ lifetime components respectively. The intermediate lifetime component resembled RAβ oligomers in buffer. In RN46A cells, this component resembled the lifetime of free rhodamine-B dye (**Fig 1B 9** and **SI 5, figure S4C**), which may indicate a possible degradation of the RAβ oligomers in these cells. The long lifetime component resembled the lifetime of RAβ oligomers in artificial lipid membranes (experimental data with artificial lipid membranes in **SI 6, figure S5**, and **table S10**). These membranes have viscosity much larger than water(27), and such an increase in rhodamine B lifetime may be expected if the dye is embedded in the membrane (28).

The most interesting lifetime component was the short one (arrow in **Fig 1B**). This was the dominant component in the astrocytes. It was also present to a significant extent in cortical neurons and in RN46A, but was absent in HeLa. Variations of environmental polarity or pH did not produce such a short component for RAβ in buffer (see **SI 7, table S11-12**), so this component likely arose from an intramolecular interaction. Since HeLa is supposed to contain less ApoE compared to brain cells, and astrocytes contain the most (19), this suggested a potential ApoE dependence of the origin of the short component. We performed immunocytochemistry (ICC) for quantifying the ApoE concentration in cells (**Fig 1C**). ICC showed the maximum intensity of the secondary antibody in Astrocytes, followed by RN46A, and then HeLa (selectivity of the primary antibody was tested, results in **SI 8, figure S6**). **Fig 1D** represents the quantification of ApoE in the different cells. Mean intensity per pixel (Alexa Fluor 647) are 32.1 ± 2.6, 17.8 ± 1.1, and 8.3 ± 0.8 (arbitrary units) for astrocytes, RN46A, and HeLa respectively. To quantify the relative fraction of the short-lifetime component in different cells, we computed the area under the short-lifetime peak normalised by the total area under the entire lifetime distribution (**Fig 1E**). Astrocytes showed the highest amount (75.2 ± 13.4%), followed by RN46A (21.2± 2.1%), cortical neurons (15.9± 2.1%), and HeLa (undetectable). While we note that ICC is not strictly quantitative, the correlation is consistent with the hypothesis that the short component is caused by the interaction of Aβ with ApoE in these cells.

To confirm the potential link between the two, we used a peptide (Aβ17-21 penta-amino acid sequence LVFFA)(29) which has been shown to block the interaction between ApoE and Aβ (synthesis and characterization in **SI 9, figure S8**). Pre-incubation of 30 µM of LVFFA in astrocytes for one hour completely abolished the short-lifetime in RAβ oligomers, leaving the other two components relatively unaffected (**Fig 1F 3**). We, therefore, conclude that the short-lifetime component provides a readout of Aβ-ApoE interaction.

### 2. Single-molecule characterization reveals changes in conformation, stoichiometry and membrane affinity of the Αβ oligomers

While FLIM is a sensitive reporter of intracellular perturbations, a detailed molecular understanding of the nature of the perturbation requires single molecule level characterizations. Our results suggested that a separate species of Aβ was formed due to the interaction with ApoE. There could be several possibilities about the formation of new species of Aβ oligomers. It could be a complex of Aβ i) with ApoE, or ii) with a cellular component whose interaction was facilitated by ApoE. It was also possible that the ApoE interaction resulted in iii) a different oligomeric stoichiometry, or iv) a different molecular conformation. Such modifications would likely result in a different size for the oligomers, which would be reflected in their diffusion constant. One of the most powerful methods to probe the difference in lifetime of the species in combination with their diffusion times is the recently developed technique of two-dimensional Fluorescence Lifetime Correlation Spectroscopy or 2DFLCS(30). 2D FLCS can distinguish different species in a mixture based on their lifetimes and sizes, and reports their interconversion dynamics with sub-microsecond time resolution(31–33). Furthermore, subsequent filtered FCS analysis can provide species associated correlation curves, hence the size (diffusion time) of each species(32,34). Also, the stoichiometry of the oligomers can be tested using single molecule photobleaching (smPB) technique(35–37).

To facilitate such detailed characterization, extracts from cells pre-treated with RAβ were subjected to 2D FLCS and smPB measurements. Extraction of the oligomers did not cause major perturbation of their intracellular state, as was confirmed by lifetime measurements (**Fig 2A**, detail of standardization of the extraction process in **SI 10**). Maximum entropy method (MEM)(38) based lifetime distribution of RΑβ oligomers (prepared freshly in buffer vs. extracted from different cells) are shown in **Fig 2B**. The conservation of the short-lifetime component post-extraction signified a cell-induced change in the RΑβ oligomers whose stability was independent of the intracellular environment. We also verified that a change of pH or the presence of divalent cations could not give rise to the short component (**SI 7, table S11-12**). We do note that post-extraction, the RΑβ oligomers showed some shift in their lifetime values (short-lifetime in RN46A extract (0.40 ns) and cortical culture extract enriched with astrocytes (0.91 ns) (**Fig 2B 3,4**). However, in both the cases, this lifetime was considerably less than that of the freshly prepared RΑβ oligomers in buffer (2.33 ns). Hence we accepted it as close to its native intracellular state. Likely, ApoE interaction did not produce a single unique species, but a distribution of new Αβ oligomers whose features were broadly conserved post-extraction.

**Figure 2.**
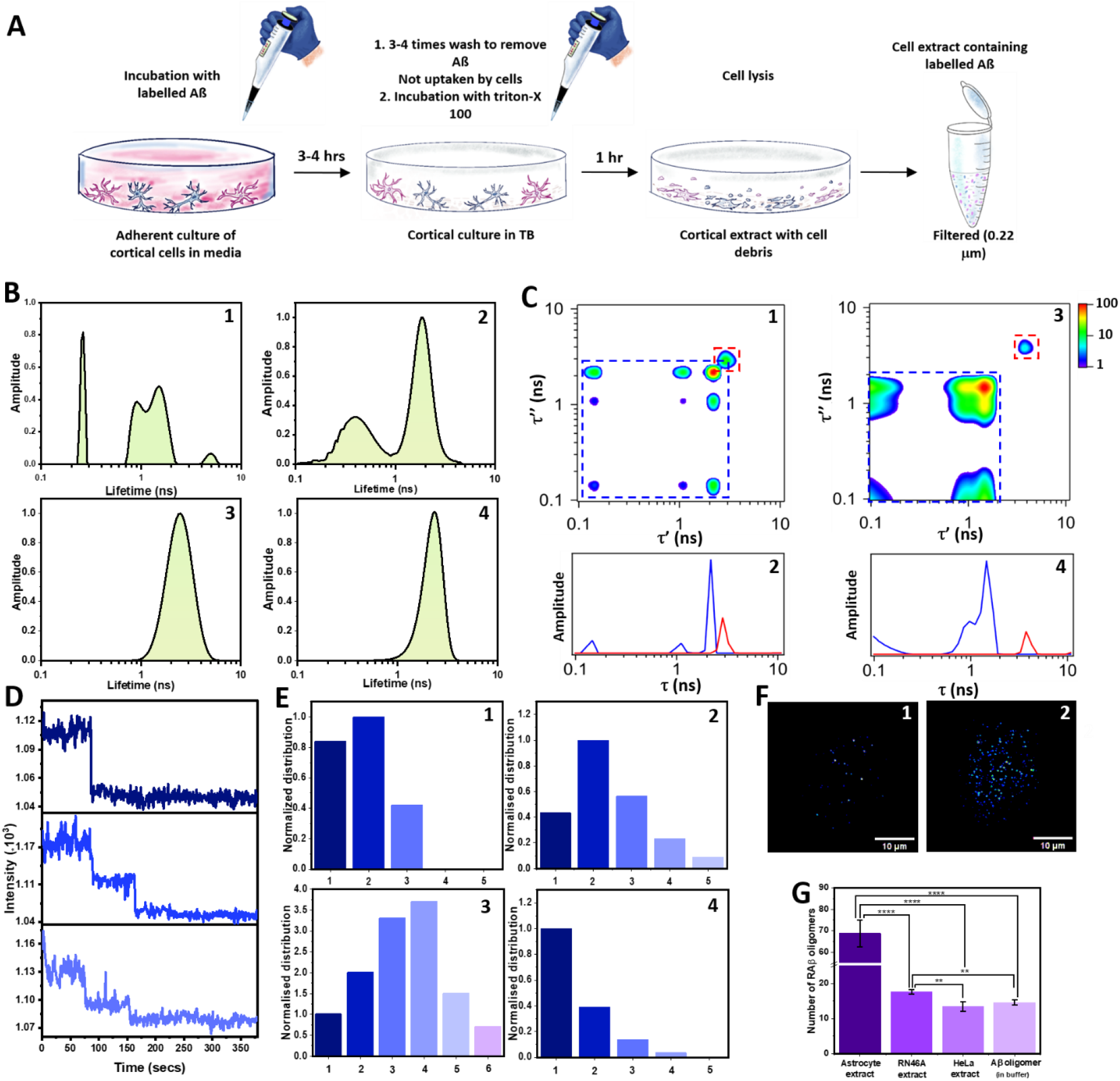
Αβ oligomer undergoes modification inside cells which are preserved in cell extracts: **A** Diagrammatic representation of the extraction procedure of RΑβ oligomers uptaken by the cortical cells. The same procedure has been used for all the different cells **B** MEM lifetime distribution of RΑβ oligomers extracted from cortical culture enriched in astrocytes (1), extracted from RN46A cells (2), extracted from HeLa cells (3), and freshly prepared in buffer (4). The persistence of the short-lifetime in the extracted RΑβ oligomers negates the possibility of intracellular stabilisation of the short-lifetime species. Thus the short-lifetime signifies a permanent modification in the RΑβ oligomers, either covalent, conformational, or both. **C** 2D FLCS probed whether the distinct oligomers are different in terms of their lifetime. 2D fluorescence lifetime correlation maps at Δ*T* = 1-20 µs (from 2D FLCS experiments) on RΑβ oligomers in buffer (1), and cortical culture enriched with astrocytes (3). The corresponding lifetime distributions of the two distinguishable species for RΑβ oligomers in buffer (2) and cortical culture enriched with astrocytes (4). Blue: short lifetime species, red: long lifetime species. The blue and red dashed squares in 1 and 3 show peaks corresponding to respective species in lifetime distribution plotted in 2 and 4. **D** Representative bleaching trajectories of individual RΑβ oligomers showing monomer, dimer and trimer. **E** Stoichiometry of the oligomers obtained from single molecule photobleaching of RΑβ oligomers extracted from cortical culture enriched with astrocytes (1), extracted from RN46A cells (2), extracted from HeLa cells (3), and freshly prepared in buffer (4). The numbers in the x-axis denote the respective stoichiometries. **F** A typical field of view under TIRF illumination of RΑβ oligomers attached to a supported lipid bilayer (PPC 111 prepared in buffer (1), and extracted from cortical culture enriched with astrocytes (2). **G** Membrane affinity of the RAβ oligomers extracted from different cells. Bar graph showing the number of RΑβ oligomer attached to the supported lipid bilayer in an area of 33.5 µm × 33.5 µm (error bars are SEM from three independent measurements). RAβ oligomers having the maximum short-lifetime shows the highest affinity to the supported lipid bilayer. The significance of difference is calculated by T-test (P < 0.0001, and P < 0.01 represented by four and three stars respectively).

RΑβ oligomers freshly prepared in buffer (with no short-lifetime component) and from extracts of cortical culture enriched with astrocytes (with major short-lifetime component evident) were used to examine whether the different lifetime components were correlated with a change in size (or diffusion constant), and whether there was a rapid interconversion between these species. **Fig 2C** shows the results of 2D FLCS on RΑβ oligomers in buffer (1-2) and in cortical culture extract enriched with astrocytes (3-4) based on a global analysis of six 2D maps. The top panels in **Fig 2C** show the earliest 2D lifetime correlation maps at Δ*T* = 1-20 µs (rest provided in SI), and the bottom panels show the fluorescence lifetime distributions of the distinguishable species. Interestingly, the RΑβ oligomers freshly prepared in buffer, which showed a single-peaked MEM distribution in **Fig 2B4**, showed two distinguishable species in 2D FLCS analysis (blue and red distributions in **Fig 2C2**). However, both these species showed intermediate mean fluorescence lifetimes (1.61±0.09 ns for blue distribution and 2.64 ±0.23 ns for red distribution) with negligible fraction of short (<1ns) components. Therefore, it is likely that the heterogeneous mixture of RΑβ oligomers in buffer, which showed a broad single-peaked MEM distribution in 1D analysis, was separated into two species with relatively close lifetime values (but both within the 1D MEM distribution) in 2D FLCS due to higher sensitivity of this technique. In comparison, RΑβ oligomers from cortical culture extract enriched with astrocytes also showed two distinguishable species in 2D FLCS analysis (**Fig 2C4**), with large differences in their mean lifetimes (1.11±0.02 ns for blue distribution and 4.18 ±0.08 ns for red distribution). Most importantly, the major species in the cell extract (blue distribution) showed a short mean fluorescence lifetime that was much shorter than both the species observed in buffer and had a lifetime distribution extending below 1 ns. This shows consistency with the intracellular FLIM data (**Fig. 1B 6**) and the lifetime fits of the extracts (**Fig. 2B 4**). Note that the distinguished species in buffer or cell extract did not show significant interconversion dynamics on the microsecond time scale (data in **SI 11b, figure S9**). The evaluation of the corresponding diffusion times of the species revealed the size of these individual species. In buffer, both the RΑβ oligomers were of similar size (diffusion times of 72 ± 15 µs and 81 ± 12 µs for blue and red respectively, data in **SI 11b, figure S10**). However, in the astrocyte extract, the two lifetimes were from two different-sized oligomers (diffusion time of 93 ± 9 µs and 280 ± 39 µs are from blue and red species respectively, data in **SI 11b, figure S10**). The longer lifetime in the cell extract aligned broadly with the long lifetime component observed in FLIM. A longer diffusion time of this species reiterated the fact that it can be lipid-associated RΑβ oligomers. This analysis also showed that the short-lifetime component did not arise from a stable ApoE-Αβ complex, as the diffusion time of such a complex would be much larger, given the size of ApoE (34 kDa). Therefore, we tested other possibilities, such as a change of oligomer stoichiometry. We used single molecule photobleaching (smPB) to measure the stoichiometry of the extracted oligomers. Stepwise bleaching trajectories of individual RΑβ oligomers (**Fig 2D**) in a single-molecule TIRF microscopy experiment reveal its stoichiometry. RΑβ oligomers prepared in buffer showed a monomer-heavy distribution (**Fig 2E 3**), whereas oligomers extracted from RN46A or cortical culture enriched with astrocytes showed a distribution with dimers as the major species (**Fig 2E 1,2**). HeLa cell extracts on the other hand showed a wider distribution with higher oligomers (**Fig 2E 4**). smPB therefore revealed a stabilization of smaller RΑβ oligomers (mostly dimers) by ApoE. These results were consistent with the small but significant increase in the diffusion times observed in 2D FLCS. Taken together, our experiments suggested that ApoE interaction caused a change in the oligomer size distribution, where dimers became the major species. We note that the lifetime distribution showed that the new ApoE-induced species was only a sub-population of the oligomeric distribution extracted from the cells. We further examined the conformation of the new oligomeric species using fluorescence quenching by a quencher (tryptophan). The results suggested that the accessibility of the N-terminal rhodamine dye was greater in the extracted species (**SI 12, figure S11, tables S15-16**), suggesting a conformation change concomitant with the change of oligomer stoichiometry.

The significant question was whether this new distribution of Αβ oligomers could be toxicogenic. To test that, we probed the membrane affinity of these oligomers using TIRF microscopy. We note that membrane interaction has been suggested as a major pathway for Aβ toxicity, and membrane affinity is a reporter of membrane interactions (39–42). **Fig 2F** shows a typical field of view of a supported lipid bilayer (detail in **SI 13b**) incubated with 500 pM RΑβ oligomers (freshly prepared in buffer and those extracted from the cortical culture enriched with astrocytes). The quantification of spot density (**Fig 2G**) demonstrated that RΑβ oligomers extracted from cortical culture enriched with astrocytes, containing a major population of the dimeric short-lifetime species, exhibited the highest membrane affinity.

### 3: Changes in Aβ correlate with its toxicity

Our membrane-affinity measurements suggested that the ApoE-induced oligomeric species might have a bearing on Aβ toxicity, and therefore we probed the toxicity of this cell-extracted Aβ species by quantifying the intracellular reactive oxygen species (ROS), a marker of cell stress(43). In neuronal cells, Αβ-induced cell stress is a well-known pathway of cell toxicity(44,45). We measured ROS together with the change of Αβ lifetime as a function of the concentration of the ApoE-Aβ interaction blocker LVFFA. The correlation between the two parameters was then tested to probe a potentially causal relationship.

As previously shown, LVFFA blocks the interaction between ApoE and Αβ oligomers, thereby preventing the development of the short-lifetime component in RΑβ oligomers. Thus, if the short-lifetime in Αβ is a reporter of its toxicogenic modification, reduction of this modification should concomitantly reduce its toxicity. **Fig 3A** shows the lifetime images of RΑβ oligomers (350 nM) in RN46A cells titrated with different concentrations of LVFFA (Lifetime distributions from each LVFFA concentration provided in **SI 14, figure S13 A**). Cell stress in RN46A induced by Αβ was measured by using the indicator dye Cell ROX deep red. Standardization of Αβ-induced reactive oxygen species (ROS) was performed (data in **SI 15, figure S14**) and 3 µM concentration of unlabelled Αβ was used for all the cell stress measurements. **Fig 3B** shows the confocal intensity images of Αβ induced ROS with different concentrations of LVFFA. Quantification of the short-lifetime component in RΑβ oligomers showed a dose dependence with increasing LVFFA concentration (**Fig 3C**). Similarly, quantification of the ROS generated by Αβ in RN46A also showed a dose dependence with increasing LVFFA concentration (**Fig 3D**). A comparison of dose dependence of both the short-lifetime and cell stress produced by Αβ is shown in **Fig 3E**. The strong correlation between the two measured parameters established that the modification in RΑβ oligomers by ApoE manifested as the short-lifetime component was toxicogenic. A non-toxic variant of Αβ (F19Cyclohexylalanine) also failed to show any short-lifetime component in RN46A cells (data in **SI 16, figure S15**). We therefore concluded that intracellular ApoE-induced modification in Αβ oligomers made them more toxic.

**Figure 3.**
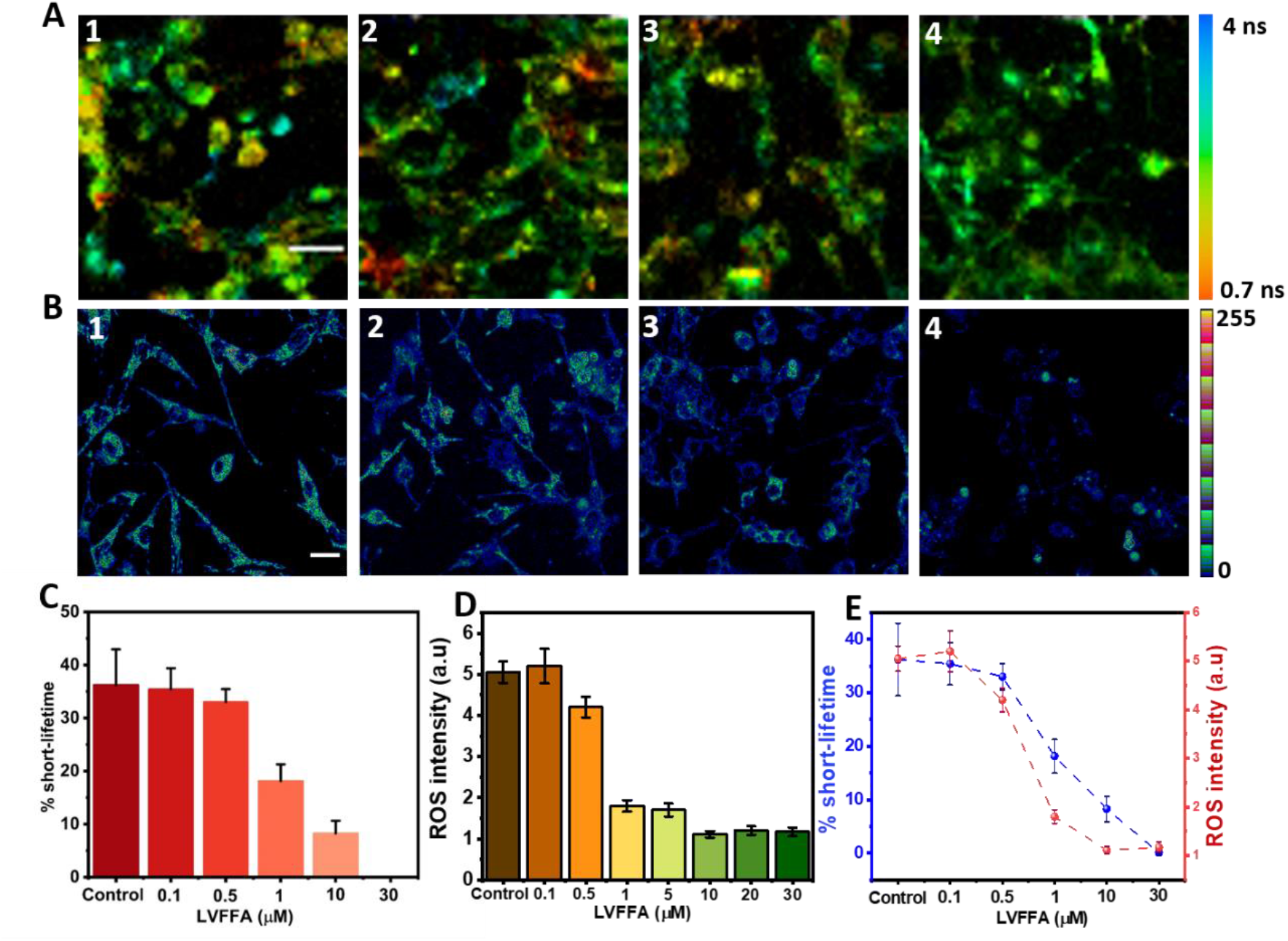
Inhibition of ApoE-Aβ interaction shows correlation between the short lifetime species and cellular stress: **A** Lifetime images of RAβ oligomers in RN46A cells, (1) 350 nM RAβ 2) 350 nM RAβ + 500 nM LVFFA, (3) 350 nM RAβ + 1 µM LVFFA, and (4) 350 nM RAβ + 10 µM LVFFA. Corresponding Aβ mediated ROS images are shown in **B**. Intensity image of 3 µM Aβ oligomers (1), 3 µM Aβ oligomers + 500 nM LVFFA (2), 3 µM Aβ oligomers + 1 µM LVFFA (3), and 3 µM Aβ oligomers + 10 µM LVFFA (4). **C** Quantification of the short-lifetime of Raβ oligomers in RN46A cells as a function of LVFFA concentration. **D** Quantification of cell stress (ROS generation) by Aβ oligomers in RN46A cells as a function of LVFFA concentration. **E** Correlation between cell ROS produced by Aβ oligomers and short lifetime of RAβ oligomers in RN46A cells as a function of LVFFA concentration. The values are connected using dashed line as a guide to the data points. (Error bars in all the measurements are SEM from three independent experiments). This clearly shows that the short-lifetime species of Aβ is more toxic than the WT Aβ oligomers. Scale bars in **A** is 40 µm, and in **B** 20 µm respectively.

### 4: Human ApoE induces a similar toxicogenic modification in Aβ oligomers in hiPSC-derived neural stem cells

Our results obtained thus far were based on experiments performed in rat brain cells (except for HeLa cells which served as a control). The ApoE protein sequence, however, is different between humans and rats(46). To probe the relevance of our findings in the context of AD, we incubated purified, recombinant human ApoE4 (artificially lipidated with DMPC vesicles) with RAβ oligomers in buffer, ensuring a near-physiological abundance ratio of 1:10 (100 nM RAβ oligomers and 1 µM ApoE)(8). To check ApoE specificity of the modification in RAβ oligomers we performed similar lifetime measurements with a few other proteins. Cytochrome C, Ubiquitin (UF45W, a tryptophan variant of ubiquitin), Myoglobin, and a standard protein BSA were used to check whether they show similar shortening of lifetime in RAβ oligomers (data in **SI 17, figure S16A**). In addition to lifetime measurements, we also checked for any complex formation between these proteins and RAβ using FCS(47). FCS showed a significant interaction with ubiquitin, in terms of increased diffusion time. Such stable complex formation was neither observed with ApoE4, nor with the other proteins (data in **SI 17, figure S16B**). On the other hand, *in-vitro* lifetime measurements of RAβ oligomers with ApoE4 showed a distinct short-lifetime, though the magnitude was less than that observed in cells (**Fig 4A**). None of the other proteins, including ubiquitin (known to form a stable complex), induced any change of lifetime. This showed that ApoE converted RAβ to a population possessing a short lifetime, but it did not remain a stable complex. These results also showed that an interaction with intracellular ApoE was much more effective in altering the lifetime. This is possibly due to the different lipidation states of the recombinant ApoE used for the *in vitro* experiments, which have been known to result in a conformational difference (10,48). A corresponding oligomer distribution of the ApoE4 incubated RAβ oligomers showed similarity with the RN46A extracted oligomers (**Fig 4B**). Our results showed that human ApoE4 could produce short lifetime Aβ oligomer species *in vitro*.

**Figure 4:**
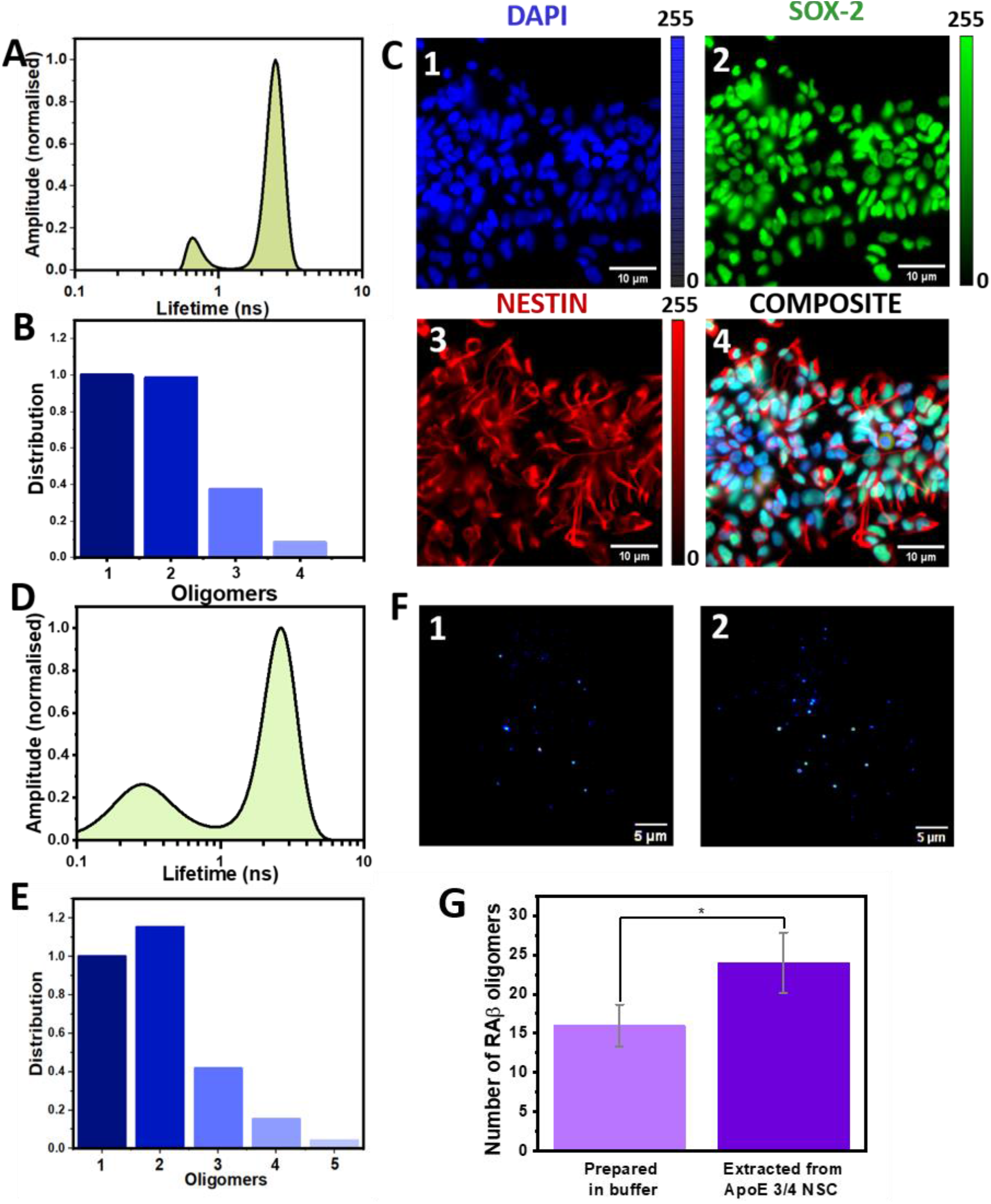
RAβ_1-40_ oligomers show short-lifetime with human ApoE isoforms both in-vitro and in cells. **A** MEM lifetime distribution of RAβ oligomers (100 nM) incubated with purified 1 µM human ApoE4 (lipidated using DMPC liposomes). Appearance of a distinct short-lifetime species can be seen. **B** Corresponding RAβ oligomer stoichiometry with ApoE4. The oligomers are different from RAβ oligomers prepared in buffer based on the origin of short-lifetime and stoichiometry. Validation of the short lifetime assay in human-derived NSCs. ApoE 3/4 NSCs were cultured to test whether the intracellular modification of RAβ with human ApoE shows any similar short lifetime. **C** Representative ICC images of iPSC-derived NSCs, stained for the nuclear marker, DAPI (1), and neural stem cell markers, SOX-2 (2) and NESTIN (3), and their corresponding composite image (4) validate the presence of neural stem cells in culture. **D** MEM lifetime distribution of RAβ oligomers extracted from NSCs (*APOE* 3/4) shows the presence of a distinct short-lifetime component. **E** Stoichiometry of RAβ oligomers in NSC extract shows a similar distribution to in-vitro ApoE modified-Aβ, with a higher proportion of dimers. Membrane affinity of RAβ oligomers extracted from *APOE* 3/4 NSC. **F** A TIRF field of view of RAβ oligomers freshly prepared (1) and extracted from the NSC culture (2) attached to PPC 111 membrane. **G** Quantification of membrane attachment shown in **F** (oligomer density from an area of 33.5 µm × 33.5 µm). Significance of difference is calculated by T-test (P < 0.05).

Further, to check the response of our lifetime-based assay in AD human samples we used neural stem cells (NSCs) generated from AD patient-derived induced pluripotent stem cells (iPSCs). NSCs, in the dentate gyrus and those derived from iPSCs, show heightened expression of *APOE*, which decreases as the stem cells mature into neurons(49,50). Moreover, the expression of different APOE isoforms in NSCs is thought to dictate the rate of neuronal differentiation and maturation(51). In this study, iPSCs were obtained from a patient with known AD background and genetic details of ApoE (Cell line No. 8904, obtained from the ADBS repository; Late-onset AD with *APOE 3/4*; detail of NSC culture and generation from iPSC in **SI 18A, figure S17**). The samples were used under appropriate ethical regulations, but these regulations did not allow FLIM imaging of the cells (which would have required transporting the cells outside the facility, so only extracts could be probed for lifetime). iPSC-derived NSCs were first stained to validate the expression of bona fide NSC marker antibodies, SOX2 and NESTIN (**Fig 4C**). The lifetime assay was performed on extracts of the NSC after RAβ oligomers were incubated (experimental protocol detailed in **SI 18B**). Lifetime measurements of RAβ oligomers extracted from the NSC showed the presence of about 5% of short lifetime (**Fig 4D**). Moreover, oligomer stoichiometric distribution showed a population similar to the RAβ oligomers extracted from rat cells (both RN46A and cortical culture enriched with astrocytes) with the dimer as the major population (**Fig 4E**), suggesting that human APOE functions in a similar manner to rat APOE, by inducing an intracellular modification of Aβ. Further, to probe whether this modification also exhibited membrane affinity-related toxicity (similar to rat cortical astrocyte extracts), we loaded NSC-derived extracts containing RAβ or freshly prepared RAβ oligomers onto an artificial supported PPC111 lipid bilayer (**Fig 4F)**. Quantification of membrane attachment (**Fig 4G**) showed that the affinity of NSC extracted RAβ oligomers was much higher than the RAβ oligomers prepared in buffer. This further supported and strengthened our findings in a disease-relevant model system, showing that APOE-induced modification of RAβ potentially perpetuates toxicity that could contribute to AD progression.

## Discussion

Studies of the molecular origin of Aβ toxicity have so far focused on their effects on cells. Studies that have employed synthetic Aβ (identical in sequence to that found in the brain) on cultured cells have mostly found the solubility and the threshold concentration necessary for observing toxic effects, to be much higher than that observed for natural Aβ in the brain (52,53). It is possible that a cellular factor interacts with intracellular Aβ, alters its conformation and oligomer stoichiometry, and enhances its toxicity. The correlation of late-onset AD with ApoE, a high-risk genetic factor, is a strong cue provided by nature, and therefore we have specifically focused on investigating potential interactions of Aβ with ApoE. We employ fluorescence lifetime of an Aβ fluorescent tag as the measurement parameter, as it is a sensitive reporter of any interaction or change of conformation near the fluorophore. Our results identify quantifiable signals in live cells which manifest such changes.

Lifetime changes of Aβ in intracellular environments had been discovered and reported by us earlier (54), but the origin of that change had so far remained obscure. So we have focused our study on different cell types containing different levels of ApoE. We find that the degree of such changes correlate strongly with the ApoE concentration in the cells (**Fig. 1**). The confirmation for this correlation arises from the well-known Aβ-ApoE interaction inhibitor LVFFA, which abolishes the particular lifetime component that bears the signature of ApoE interaction (**Fig. 1F3**). Specific molecular level changes underlie this lifetime perturbation, as shown in our studies with the Aβ cell extracts. The major question is whether ApoE makes a complex with Aβ to induce the lifetime changes. 2DFLCS correlates the lifetime of a chemical species with its size. It shows that while a fraction of the Aβ binds something much larger, the short lifetime component originates not from this species, but from species that is only about 20% larger than Aβ. The smPB results show that these changes in the diffusion time are most likely due to an increase in the dimeric population of the oligomers compared to the monomer-rich population observed for Aβ in buffer.

Further, to test if these changes are toxicogenic, we probe intracellular ROS activity in rat RN46A cells. The use of LVFFA at different concentrations show a clear correlation between the short lifetime component and the cellular stress as measured by ROS concentration. The correlation is strong, but the two graphs do not coincide. However, toxicity, or cellular stress, is likely to have threshold-sensitive effects, so a linear correlation between the two may not be expected.

Finally, we probe whether these findings are relevant to human AD-specific model systems. We examine the emergence of a short lifetime component when RAβ is incubated with human ApoE4 *in vitro*, and found a small but significant component associated with it. While the relatively low strength of this component can have its origin in the different lipidation state of ApoE, the uniqueness of this effect signifies that human ApoE also similarly interacts with Aβ. We further corroborated this by investigating this effect in human neuronal stem cells (ApoE3/4), and find a very strong correlation of the effect observed already in rodent cells. The oligomeric distribution and the affinity for lipid bilayer membranes also changes in accordance with this modification (as was also shown by rat ApoE), indicating that human ApoE causes a similar effect. This suggests that APOE causes a modification in the oligomeric state of AB, resulting in an increase in toxicity.

In summary, the change in the N-terminal region of Aβ oligomers is reflected in the fluorescence lifetime of a tag and shows a positive correlation with the concentration of ApoE. These changes can be rescued with a known inhibitor of ApoE-Aβ interactions. This results in the increase of oligomers of a specific size, which have a higher affinity to lipid bilayers and cause oxidative stress in cells. These results are reproducible in human-derived neuronal stem cells.

Our study thus offers proof of principle which shows that ApoE causes a transition of Aβ oligomers to a more toxic state and that this can be directly imaged in living cells. This provides a foundation for future studies in understanding the molecular aetiology of AD and for the development of therapeutic strategies.

## Acknowledgments

We acknowledge generous help from Dayana Surendran, Sushanta Chhatar, Aditya Shrivastava, Sudip Pal and Sreelakshmi C at different stages of this project.

## Funding

Accelerator program for Discovery in Brain disorders using Stem cells (ADBS) seed grant, Dept. of Biotechnology, Government of India, no. 8251-5 (OM, SM)

Department of Atomic Energy, Government of India, project no. RTI4003 (SM)

## Author contributions

Conceptualization: AD, OM, SM

Methodology: AD, BS, KG, KI, TT, OM, SM

Investigation: AD, AV, AKD, UB, BS, VV, MK

Funding acquisition: OM, SM

Project administration: SM

Supervision: KG, KI, TT, OM, SM

Writing – original draft: AD, BS, SM

Writing – review & editing: AD, KI, TT, AV, AKD, OM, SM

## Competing interests

Authors declare that they have no competing interests.

## Data and materials availability

All data will be made available upon request. Material except hiPSC will also be available upon request.

## Notes

### Competing Interest Statement

The authors have declared no competing interest.

