## Supporting information for "Apolipoprotein-E transforms intracellular Amyloid-β oligomers to a more toxic state"

### Summary

The supporting information is divided into materials and methods, supplementary figures, and supplementary tables accordingly.

### Materials and Methods

#### SI 1: Primary dissection of Sprague Dawley pups and mammalian cell culture

Dissection of WT Sprague Dawley pups was performed to isolate the cortex. This acted as a source of astrocytes, and cortical neurons for our study. The dissection was followed using a standard procedure as follows.

a) Pups (P0-P4) were anesthetized by keeping them in ice for approximately 5 minutes.
b) 60 mm petri dishes were kept ready with ice cold HHGN buffer or Thomson's buffer (composition mentioned below)
c) Pups were taken out of the ice and sprayed with 70% ethanol. Decapitation was performed and the head was transferred on to one of the dishes.
   - Under a dissection microscope, the skin was removed with a fine scissor. For removing the skull, the eyes were held with forceps and the skull flaps were removed. This makes way to the brain which was transferred to the second petri dish.
   - The two brain hemispheres were separated. Removal of the olfactory lobe facilitated the removal of the meninges which was pulled in a single movement while opening the hemisphere.
   - Separation of the hemispheres exposed the hippocampus and the cortex. The hippocampus was removed from the cortex to obtain a pure cortical culture.
   - Cortex was cut into small pieces, and collected in a tube with HHGN/ Thomson's buffer and stored in ice.
d) Once the cortices are collected, they were allowed to settle at the bottom of the tube. The buffer was removed and interchanged with fresh buffer containing 1X trypsin-EDTA.

e) The solution was incubated at 37°C for 15 minutes, agitated, and re-incubated for another 15 minutes.

f) After 30 mins, the trypsin solution was removed, followed by three buffer washes. Ice cold buffer was added and triturated till a homogenous solution is obtained.

g) The cell suspension was centrifuged at 700-800 RPM for 5 mins to pellet the cells down.

h) Finally, the buffer was replaced with pre-heated media (37°C, composition provided below) and cells were plated in required density in flasks (T-25 or T-75) and incubated at 37°C.

i) Half media change was performed after 48 hours, and full media change after 7 days. Afterwards, trypsinization and regular media change ensured enriched astrocyte culture. For obtaining cortical neurons, suspension obtained from first trypsinization was plated directly at low density and trypsinization was avoided.The entire steps followed in dissection is shown in figure S1.

High passage (p) cells of RN46A (rat serotonergic neuron-derived cell line), and HeLa (P-2,3) cells were cultured in DMEM-F 12 (1:1) (Gibco, USA) media supplemented with 5% FBS, 50 units/ml Penicillin and 50 µg/ml Streptomycin (Gibco, USA) at 37˚ C (for growth). Poly-l-lysine coated T-25 canted neck flasks (Falcon, USA) were used for growing the cells in humidified air containing 5% CO2. Media was changed every 48 hours. For RN46A cells, G418 supplement was added at a concentration of 50For imaging or FLIM studies, cells (at least two passage after fresh thawing) were grown on poly l lysine2 coated 0.22 um coverslips.

**SI 2: Identification of astrocytes in culture**

To identify the presence of astrocytes in the culture, standard immunocytochemistry (ICC) was done. The procedure of ICC remains the same in all the cases except specific antibodies in each cases. For identifying the presence of astrocytes in the cortical culture, cells were plated in square coverslips (pre-coated with 100 µg/ml of poly-l-lysine). Briefly, the procedure is outlined as follows

a) Adherent cells were washed gently with warm PBS buffer (pre-heated at 37°C) three times to remove loosely attached cells and media
b) Thereafter, the cells were fixed using 2% paraformaldehyde (PFA) for 20 minutes and washed with PBS 2 times. The incubation was carried out at room temperature (RT).
c) Excess PFA was quenched by using a PBS solution of 0.1 M Glycine. Incubation was carried out at RT for 15 minutes followed by 2 times washing using PBS.
d) To introduce the antibodies, cells were permeabilized using 0.3% Triton-X 100 (in PBS) for 5 minutes.
e) Removal of excess Triton X was carried out by washing the cells 2 times using PBS for 2 minutes each under shaking.
f) Blocking was performed with 3% bovine serum albumin (BSA) in PBS for 45 minutes at RT followed by a single quick PBS wash.
g) Rabbit polyclonal anti-glial fibrillary acidic protein (GFAP) antibody (G9269, Sigma Aldrich, reactive with rat and human) was used as a primary antibody. A dilution of 1:500 by volume (antibody:PBS) was used to stain the cells. Incubation was carried out at RT for 4 hours.
h) Post incubation, cells were washed 3 times for 5 minutes each under shaking. Goat anti-rabbit IgG Atto 488 antibody (18772, Sigma Aldrich) was used as a secondary antibody. A volume dilution of 1:1000 (antibody:PBS) was used to stain the cells for 1 hour at RT.
i) Cells were washed thrice for 5 minutes each under shaking. Post washing, the cells were mounted using mounting media in coverslides and were either stored at 4°C or imaged immediately.

Subsequently, the shape of the cells was used to recognize the presence of astrocytes for fluorescence imaging. Identification of the astrocytes using GFAP is provided in figure S2.

### SI 3: Experimental outline for intracellular lifetime measurements of RAβ oligomers

To check whether intracellular ApoE content and lifetime modification (short-lifetime component) in RAβ oligomers are correlated, we performed FLIM imaging of RAβ oligomers in different cells of different content. Cells (primary rat pup astrocytes, cortical neurons, RN46A, and HeLa) were incubated with RAβ oligomers (350 – 500 nM) for 3 hours in cell media. RAβ oligomers were aliquoted, flash frozen, and stored at pH 11 at high concentration to prevent aggregation. These aliquots were either directly diluted in buffer or incubated in cells at appropriate concentration. Post incubation, cells were washed with Thomson's buffer (TB) and different region of Interests (ROIs) were imaged in TB. Pixel wise lifetime decay in each image was fitted with two exponentials in SPCImage analysis software (Becker and Hickl). Average lifetime was calculated using a MATLAB code (written in lab).

### SI 4: Composition of HHGN buffer, Thomson's buffer, and primary culture media

For dissection of rat pups and subsequent washing, either HHGN buffer or Thomson's buffer was used. The composition of HHGN buffer (500 ml) is provided in table S1.

In the preparation of the HHGN buffer, another buffer composition, HBSS is required. Thus, the composition of HBSS buffer (10X, 100 ml, pH 7.5) is provided in table S2. Preparation of Thomson's buffer (T.B, 1X, 50 ml) is provided in table S3. In each case, the prepared buffer is filter sterilized with 0.22 µm filter prior to usage.

The disintegrated cells from the cortex were cultured in primary media. Preparation of 100 ml of primary media (and its composition) is provided in table S4.

### SI 5: Fluorescence Lifetime imaging and calculation of the short-lifetime component of RAβ oligomers in cells

The cells were incubated with the N-terminal rhodamine B labelled Aβ oligomers (RAβ) for 3 hrs in the cultured media at 37° C. After they were washed with TB, were subjected to FLIM experiments. For FLIM measurements, a super continuum pulsed white light laser (repetition rate was 80 MHz, but could be reduced by a factor of 2, 4, 8, 16 or 32, as required) was used as the excitation source. A light beam of 540 nm (± 3 nm) wavelength was separated from the white light laser source using an acousto-optics filter to excite the Rhodamine-B molecule. The incoming laser beam was expanded using a telescope set up, and the expanded beam is directed towards the back aperture of a 1.2 NA 60x objective (Olympus America Inc, Center Valley, PA, USA) using a dichroic mirror (552 nm long pass; Chroma, VT, USA). The objective lens focused the laser light on the sample placed on a coverslip. The fluorescence was collected back using the same objective lens, separated from the excitation by a dichroic mirror, and then focused into a fibre coupled single photon avalanche photodiode with high time resolution (SPAD, Micro Photon Devices, Italy). The core diameter of the fiber was 50 µm. The instrument response function (IRF) was recorded using a sample consisting of a mixture of chalk powder and bleaching powder. During the IRF measurement, a 530/43 filter was used just before the optical fibre to reduce scattered light intensity. The full width half maxima (FWHM) of the response function was ~ 160 ps. While collecting fluorescence from sample, a 607/73 emission filter was used to block any backscattered light. The fluorescence was collected at different points of the sample by scanning the laser beam in the XY plane. The scanning was achieved by pixel-wise raster scanning of the galvanic mirrors of the confocal microscope. TCSPC data was collected from each pixel and then analysed for the fluorescence lifetime to produce the lifetime image. This required the TCSPC acquisition to be synchronized with the pixel clocks of the scanning mirror, which was achieved through a cable connecting the laser scanning microscope (LSM710) with SPC150 TCSPC data acquisition card. The analysis was done using SPC Image software.

The pixel-wise lifetime decays from each cells were fitted using two exponential decay components. Average lifetime distribution was calculated as,

$$\tau_{avg} = \frac{A_1\,{\tau_1}^2 + A_2\,{\tau_2}^2}{A_1\,\tau_1 + A_2\,\tau_2}$$

A distribution of the average lifetime shows three distinct populations of RAβ oligomers. These were fitted independently using 3 Gaussians. The fits are provided in figure S3.

From these individual lifetime distributions obtained, the percentage contribution from each component was calculated by their relative amplitude (that is, the contribution of a particular component was calculated as area under that Gaussian divided by the total area under the curve). Thus, the three corresponding lifetimes and their contributions in each cell types are provided in table S5.

From these values, the average lifetimes were calculated for global fitting. The first population was assigned a lifetime of 530 ps, the second and third populations as 1974 ps and 3270 ps respectively. For global fitting, a range of 20% was allowed for each of the lifetime components to fit the distributions from all the cells. In case of HeLa, individual fit was unable to fit with 3 components, thus the distribution was fitted with the two Gaussian components using the same parameters from the Global fit. The global fit parameters are provided as in table S6, S7, S8, and S9.

The distinct average lifetime populations so obtained were classified into a short lifetime component, long lifetime component, and a component that resembled freshly prepared RAβ oligomers in TB (provided in the main text, lifetime around 2.3 ns) or that of rhodamine B in cells (particularly in case of RN46A).

The lifetime population of RAβ oligomers less than free rhodamine B in cells is attributed to a short-lifetime component. This points towards an intracellular dynamic quenching in the N-terminus label which could be either because of a conformational change in RAβ oligomers (its own amino acids tyrosine, arginine can show dynamic quenching with rhodamine B**,** figure S4A, B) or some intracellular interaction. Lifetime of free rhodamine-B in RN46A cells shows 2 populations around 1.8 ns and 2 ns respectively (figure S4C, D).
As can be seen from the average lifetime distribution of RAβ oligomers in RN46A cells, one of the lifetime component resembles that of rhodamine B in RN46A cells. This indicates some degradation of RAβ oligomers particularly in RN46A cells.

The longer lifetime component can be attributed to RAβ oligomers attached to lipid membrane. *In-vitro* measurement of the lifetime of RAβ oligomers in PPC -111 small unilamellar vesicles (SUVs) shows a longer lifetime.

### SI 6: Lifetime of freshly prepared RAβ oligomers in membrane

To determine the lifetime of the RAβ oligomers in membrane, we prepared small unilamellar vesicles (SUVs) of POPC:POPG:Cholesterol (PPC 111) lipid via sonication (in TB) and incubated freshly prepared RAβ oligomers (350 nM) with the vesicles (2.5 mg/ml) for 60 minutes. We have shown previously that RAβ oligomers strongly binds to small vesicles[1]. Thus a lifetime measurement of the solution would be dominated by the lifetime of RAβ oligomers in membrane. TCSPC decays of freshly prepared and membrane incubated RAβ oligomers are given in figure S5. Table S10 contains the fit parameters and the average lifetime values. It can be observed that the average lifetime of the RAβ oligomers are longer in membranes.

### SI 7: Effect of pH and EDTA on the short-lifetime component of RAβ oligomers extracted from RN46A cells

As several divalent metal ions such as $Zn^{2+}$, $Cu^{2+}$ are known modulators of Aβ toxicity[2,3], we checked whether the origin of the short-lifetime in cells is because of some metal ion. Thus the extracted RAβ oligomers (from RN46A cells showing the short-lifetime) was treated with 20 mm EDTA. The short-lifetime persisted the EDTA incubation hinting to the fact that the origin of short-lifetime was not because of any metal ion.

Similarly, we modified the pH of the cell extract to more acidic (pH 4.5) to check whether any ionic interaction is involved in the origin of the short-lifetime component. The change in Ph of the buffer also did not abolish the short-lifetime component. Thus it suggested it was neither because of any ionic interaction as well. Tables S11, and S12 contain the lifetime decay parameters for the treatments of cell extracted RAβ oligomers with different pH and EDTA.

### SI 8: Intracellular ApoE quantification (western-blotting of ApoE antibody and ICC)

Intracellular ApoE was quantified using Immunocytochemistry. Prior to that, the primary antibody used (abcam, EPR19392) to detect ApoE was tested for its specificity by Western blotting using WT Sprague Dawley pups brain lysate and control HeLa. Briefly, the western blotting procedure is mentioned below.

**Cell Culture and Harvesting**

a) Isolated primary cortex from the rat brain (Sprague Dawley pups P0-P4) was flash frozen and stored at -80°C. HeLa cells were grown in T-75 flask to a confluency of 80-90% for preparing cell lysate.

b) Ready to harvest tissues were weighed. 100 mg of tissue was added to 500 μl mixture of RIPA lysis/extraction buffer (thermo Scientific, 89901) and protease inhibitor tablets (1X, Roche, 04 693 159 001). Tissues were grind finely in homogenizer and kept in ice for 30 mins. For HeLa cells, 500 μl of the same cocktail was added to the flask and the cells were scraped. The cell lysate was also kept in ice for 30 minutes.

c) Post incubation, the lysates were centrifuged at 12,000 RPM for 30 minutes T 4°C to clear the cell debris.

d) The supernatant from each sample was collected. This was the total protein sample used for further estimation.

**Protein Quantification**

a) The total protein concentration both in the cortex and HeLa extract were estimated using a BCA assay kit (thermos Scientific, 23225) standardized with known concentrations of BSA. The obtained concentration of total protein was 4.3 mg/ml, and 4.4 mg/ml for Cortex and HeLa respectively.

b) This was important to load same amount of total proteins in the gel.

**SDS-PAGE**

a) Sample preparation - Protein samples were prepared by loading Laemmli buffer (containing β-mercaptoethanol, glycerol, Tris-Hcl, SDS, and bromophenol blue) to the protein sample in a ratio of 1:3 (dye buffer:sample). Samples were heated at 95°C for 8 minutes to denature the proteins. The samples were cooled and used immediately.

b) Gel preparation - For performing the SDS-PAGE, both loading gel and resolving gels were prepared. The composition of both are provided in tables S13, and 14. For ApoE, we prepared 10% polyacrylamide gel.

c) Running buffer was prepared (1 Lt buffer contains Tris base 3 gm, Glycine 18.75 gm, and SDS 1 gm, pH 8.3). Equal concentration of proteins for the two samples were added to the lanes of the gel.

d) For docking (initial 10-15 minutes), the gel was run at 80 V (Bio Rad vertical gel electrophoresis system). After that, the entire run was completed at 130 V until the dye front reached the bottom of the gel.

**Protein Transfer**

a) Protein bands in the gel were transferred to a PVDF membrane using a Bio-Rad mini trans blot (1703812). The membrane was activated prior to use in methanol for 5-10 minutes.

b) Transfer buffer (1 Lt contains Tris 3.025 gm, glycine – 14.26 gm, and methanol 200 ml) was prepared to carry out the entire transfer process.

c) The entire transfer was carried out in cold room (4°C) at 300 V for 2 hours.

**Blocking and Antibody Incubation**

a) Once the transfer is done, the membrane was washed thoroughly with transfer buffer. The front and back ends of the membrane was marked. Blocking was performed with 5% BSA solution (30 minutes rocking, RT).

b) Primary antibody was prepared (1 µl in 2 ml transfer buffer). The blocking solution was discarded from the membrane and the primary antibody was added. No in-between washing was performed.

c) The primary antibody was incubated overnight at 4°C in a rocker.

d) Post primary antibody treatment, the membrane was thoroughly washed with transfer buffer and Tween-20 (Sigma-Aldrich, 9005-64-5) 5 times, 5 minutes each. Tween-20 prevents the membrane from drying, but at the same time prevents antibody attachment. Thus, a final wash in normal transfer buffer withot Tween-20 was performed.

e) A secondary HRP-linked antibody (0.75:1000 µl in Transfer buffer, cell signalling technologies, 7074S) was used to stain the membrane for 30 minutes (rocking at room temperature)

f) Post incubation, the membrane was washed with transfer buffer (containing Tween-20) for 5 times, 5 minutes each.

g) The membrane was ready to be developed.

**Membrane development**

a) Membrane was developed in a Syngene G:Box iChemi XR Gel Documentation and Analysis System (GelDoc, Gel Imager).

b) Chemiluminescence of the HRP was used to detect the bands. No observable band was seen for HeLa extract whereas a clear band around 36 kDa could be seen in the brain cortex extract.

**Loading control**

To measure the loading control, the same membrane was stripped and amount of GAPDH (loading control) was checked using a primary antibody (2118L, cell signalling technologies) and the same secondary antibody.

In figure S6, the bands in the blot were not intensity quantified as a clear ApoE band was seen in the cortex extract lane and no observable bands were observed in the HeLa extract lane. Having checked the specificity of the anti-ApoE antibody, ICC was used to quantify the amount of ApoE in the cells (figure provided in main text). As a control, all the ICC's were repeated in the same cells without the primary antibody to detect the presence of any undesirable background from the secondary antibody. Figure S7 clearly shows there are no non-specific backgrounds coming from the cells. This also helps to correlate the quantify the ICC results for ApoE reliably with the intracellular ApoE content.

**SI 9: Synthesis, purification and incubation of penta-amino acid peptide LVFFA**

Synthesis of the penta-amino acid peptide LVFFA was performed using standard Fmoc chemistry in an automated solid phase peptide synthesizer (PS3, Protein Technologies Inc., USA). Rink amide MBHA resin LL (0.31 mmol/g) was used as a solid support to grow the peptide chains. 4-fold excess Fmoc protected amino acids were activated with equimolar HATU or HBTU and NMM (0.4 M) in DMF. After that, Fmoc was deprotected using mixture of 2% DBU and 20% piperidine in DMF (v/v). A mixture containing TFA, TIS, water, and phenol at a volume ratio of 85:5:5:5 was used for the cleavage of the peptides from the resin and deprotection of the acid labile side chains (mixing time 4 hours). The peptide was concentrated under nitrogen flow, then precipitated and washed with tert-butyl methyl ether. The precipitates were dried under vacuum to obtain powdered. Obtained powder was dissolved in 50% ACN: water mixture and purified by reverse phase high performance liquid chromatography (Prominence 20A, Shimadzu Corporation, Kyoto, Japan). Separation of dissolved species was achieved by passing the crude peptide through a C16 analytical column (Kromasil, Eka Chemicals AB, Bohus, Sweden) and using a gradient of acetonitrile and 0.1% TFA in water as the eluent. The purity of the peptide was tested by MALDI

(Model: TOF SPEC 2E, Micromass, Manchester, England). The purified peptides were lyophilized and stored at -80 ºC under desiccated condition until used.

A matrix assisted laser desorption/ionization (MALDI – TOF) mass spectra of the peptide synthesized is provided in figure S8. (The expected mass of the LVFFA peptide is 594.3. We get a sodium adduct 617.2 (m/z = (594.2 + $Na^+$)/1) and a potassium adduct 633.2 (m/z = (594.2 + $K^+$)/1).

**SI 10: Protocol to isolate intracellular RAβ oligomers from cells**

To isolate the intracellular RAβ oligomers from different cells, cells were incubated with 350-500 nM RAβ oligomers for 3 hours. Previously we have established that incubation of RAβ oligomers in cells under identical experimental condition gives rise to a short-lifetime. Thus, cells in T-25 flasks were incubated with RAβ oligomers for 3 hours. Post incubation, cells were washed off excess unattached RAβ oligomers with TB thrice. Triton-X 100 in TB (100 µM) was added to the cells and the entire system was kept in 4°C for 1 hour. Post incubation of the surfactant, the entire cell suspension was filtered using a 0.22 µm PVDF syringe filter. The filtered suspension was checked for rhodamine B fluorescence and lifetime measurement. Appearance of a short-lifetime component in the suspension ensured isolation of the native lifetime modified RAβ oligomers in buffer.

**SI 11a: 2D FLCS - Experimental details**

The 2D FLCS measurements are performed on ~10 nm R□□□-40 in buffer (PBS: 10 mM $Na_2HPO_4$, 137 mM NaCl, 2.7 mM KCl, pH 7.4) or astrocyte extract using a custom-built time-correlated single-photon counting (TCSPC)-fluorescence correlation spectroscopy (FCS) set up following established protocols [Ishii and Tahara 2013, Sarkar et al JPCL 2019]. Briefly, a supercontinuum laser operating at 30 MHz (Leukos Rock-400-4-PP, France) is used as a pulsed excitation source. The broadband light generated by this laser is filtered using a bandpass filter (Semrock FF01-525/30, USA) and introduced into a confocal microscope (NIKON Eclipse Ti, Japan) using a single-mode optical fiber (Thorlabs P5-460B-PCAPC-1, USA). In the microscope, the filtered excitation light is collimated using an achromatic lens and focussed onto the sample using a water immersion objective lens (Nikon Plan Apo IR 60×, numerical aperture: 1.27). We set the power of the excitation light at 50 μW at the back aperture of the objective lens. The sample chamber is assembled using two coverslips separated by a silicone spacer. The fluorescence from freely diffusing sample is collected by the same objective lens, separated from excitation light using a dichroic mirror (Chroma ZT532/640rpc, USA), and guided into a multimode optical fiber (Thorlabs M50L02S-A, USA). The entrance of this multimode fiber (50 μm core diameter) acts as the confocal pinhole for this set up. The multimode fiber directs the collected fluorescence light into a detection assembly (Thorlabs DFM1/M, USA), where the fluorescence is filtered using a dichroic mirror (Chroma ZT633rdc, USA) and a bandpass filter (Chroma ET585/65m, USA). Subsequently, individual fluorescence photons are detected using a hybrid detector (Becker and Hickl HPM-100-40-C, Germany) and recorded using a TCSPC module (Becker and Hickl SPC-130EM, Germany) operating in the time-tagging mode. The TCSPC is synchronized with the pulsed laser excitation by detecting a part of the excitation light using a fast photodiode (Positive light).

For each detected photon, we recorded its macrotime, $T$ (arrival time from the start of the experiment) and microtime $t$, (arrival time from previous excitation pulse). For 2D FLCS[4,5], we utilized this information of $T$ and $t$ of all detected photons to construct six 2D emission decay maps at macrotime intervals $\Delta T$ = 1-20 μs, 20-50 μs, 50-100 μs, 200-500 μs, 1-2 ms, and 3-5 ms. These six 2D maps are then globally analyzed with 2D inverse Laplace transformation using maximum entropy method (global 2D MEM analysis) to obtain corresponding 2D lifetime correlation maps, as well as the fluorescence lifetime distributions of the distinguished species and the correlations between them at each $\Delta T$. Mathematically, a 2D lifetime correlation map at $\Delta T$ is represented as

$$\widetilde{M}(\Delta T;\tau',\tau'') = \sum_i^{n_{sp}} \sum_j^{n_{sp}} a_i(\tau') g_{ij}(\Delta T) a_j(\tau''),$$

where $n_{sp}$ is the number of distinguishable fluorescence species, $a_i\ (\tau)$ is the fluorescence lifetime distribution of species $i$, and $g_{ij}(\Delta T)$ is the correlation between species $i$ and $j$ at $\Delta T$. Here, $i = j$ indicates autocorrelation and $i \neq j$ indicates cross-correlation. Based on these correlation values, we determined the conversion ratio between species *i* and *j* at Δ*T* as

$$R_{ij}(\Delta T) = \frac{g_{ij}(\Delta T)}{\sqrt{g_{ii}(\Delta T) \times g_{jj}(\Delta T)}}.$$

R(Δ*T*) varies between 0 and 1, where 0 indicates no interconversion dynamics and 1 indicates equilibrium. For filtered FCS analysis[6,7], we determined the fluorescence decay of each distinguished species by performing Laplace transformation on corresponding fluorescence lifetime distribution, and then utilized them to determine species-specific auto and cross-correlation functions. For determination of the diffusion time of each species, we fitted its autocorrelation function for a single freely diffusing species in 3D with contributions coming from triplet state formation of the dye. All analyses are performed using custom written programs on Igor Pro (Wavemetrics, USA).

**SI 11b: 2D FLCS - characterization and comparison of RAβ oligomers (freshly prepared and extracted from cortical culture)**

By determining the correlations between the major and minor species across six 2D maps (Figure S9A, B and Figure 2C in main text), we observe that the two species do not show significant interconversion dynamics on the microsecond time scale for either sample (Figure S9C).

To glean more information on the major and minor fluorescence species in each sample, e.g., their relative population, diffusion times, and interconversion dynamics, we determined individual autocorrelations of these species and the cross-correlations between them by performing filtered FCS analysis using their fluorescence decays (obtained with Laplace transformation of the fluorescence lifetime distributions in **figure S9B**)[6,7]. Based on the initial autocorrelation amplitudes of the two species, the population of the major species is determined to be 5.6 times that of the minor species in astrocyte extract, whereas it is determined to be 2.3 times in buffer (**figure S10 A, B**). By comparing their diffusion times, the major species in buffer likely represents RAβ monomers, whereas the minor species, showing ~10% longer diffusion time, likely represents RAβ oligomeric mixture rich in dimers. These results are consistent with those observed in smPB experiments (**figure 2E 3,** main text) and indicate a monomer dominated mixture of RAβ monomers and small oligomers in buffer. In comparison, the diffusion time of the major, short fluorescence lifetime species in astrocyte extract is ~10-20% larger than that in buffer, likely indicating a RAβ oligomeric species (consistent with **figure 2E 2,** main text). Interestingly, the minor, long fluorescence lifetime in astrocyte extract shows ~3 fold longer diffusion time compared to the major species. This slow diffusion possibly comes from binding of RAβ to cellular component, e.g., lipid membrane. This binding also likely leads to an increase of fluorescence lifetime of this minor species to 4.2 ns. We notice that the cross-correlation amplitudes between the major and minor species are very low in both samples (**figure S10C, D**). Consistent with the conversion ratio curves obtained in 2D FLCS analysis, these low cross-correlation amplitudes also indicate that the two species do not undergo significant interconversion on the microsecond time scale. In fact, we can also determine a continuous conversion ratio curve across all $\Delta T$s by normalizing these cross-correlation curves with the autocorrelation curves of the major and minor species (**figure S10E**). We note that these curves show a weak rise on the millisecond time scale in both samples, indicating the possibility of interconversion dynamics on the millisecond time scale. However, this rise cannot be confirmed in the present data because of poor S/N ratio on the millisecond time scale as a result of molecular diffusion on faster time scales.

#### SI 12: Dynamic quenching of RN46A cells extracted RAβ oligomers

To check whether the RAβ oligomers of different lifetimes have different conformation, we performed dynamic quenching of the RN46A extract. Since the extract contains both short-lifetime and long lifetime species, different quenching efficiency by a quencher would indicate differential N-terminus accessibility of the two types of RAβ oligomer. Tryptophan quenches fluorescence of rhodamine B by both dynamic quenching and complex formation[10]. Thus, we titrated the cell extracted oligomers with increasing concentration and measured the lifetime changes of the respective species. Figure S11 contains the lifetime decays, and tables S15, and 16 contains all the fitted parameters for tryptophan quenching of RAβ oligomer extracted from RN46A culture and freshly prepared in buffer.

**SI 13a: Single molecule photobleaching (smPB) – experimental details**

To perform smPB of RAβ oligomers (derived from different cell extracts or freshly prepared in TB), we used a home-built objective based total internal reflection fluorescence (TIRF) microscope [8]. Briefly, a 543-nm He-Ne (25-LGR-393-230; Melles Griot, Rochester, NY) laser was used for focused excitation (back aperture power 1.3 mW) at the back focal plane of a Nikon APO TIRF 100X/1.49 objective. A dichroic (565 nm) was used to separate the fluorescence from the excitation. The fluorescence was collected using a band-pass filter (605/55 nm, BA577-633, Nikon) and focused onto an electron multiplying CCD camera (ANDOR iXON, DV887ECS-UVB) using a 50 cm biconvex lens. A series of frames were captured (frame rate 90 ms) until the fluorescence spots in a given location were almost fully bleached. For identifying stoichiometry of the spots and determining the step lengths. Individual spots from this dataset were analysed for identifying the stoichiometry. A diagrammatic of the home-built TIRF setup is provided in figure S12.

**SI 13b: Sample preparation to identify the stoichiometry and quantify membrane attachment of RAβ oligomers**

In smPB to detect the stoichiometry of fluorescent multimers, the fluorescent markers should be stationary. This allows correct monitoring of the bleaching steps. Hence, to immobilize the fluorescent spots, either samples were spin coated or attached to / distributed on a lipid bilayer. For spin coating, samples (freshly prepared or cell extracted RAβ oligomers) were diluted in 0.25% poly vinyl alcohol (PVA) solution to prepare a final concentration of 0.5-1 nM. On precleaned coverslips (both piranha and plasma treated), PVA sample solution was spin coated (3000 RPM) for 30 seconds. This ensured uniform coating and well dispersed fluorescent spot density for imaging.

For measuring membrane attachment of RAβ oligomers, supported lipid bilayer (SLB) were prepared. For preparing PPC 1:1:1 bilayer (POPC: POPG: Cholesterol in the molar ratio 1:1:1), the lipids 1-palmitoyl-2-oleoyl-sn-glycero-3-phosphocholine (16:0–18:1, PC) (POPC), 1-palmitoyl-2-oleoyl-sn-glycero-3-[phospho-rac-(1-glycerol)] (16:0–18:1 PG) (POPG) and cholesterol were purchased from Avanti Polar Lipids

(Alabaster, AL) and from Sigma-Aldrich (St Louis, MO) respectively. SLBs were prepared using vesicle fusion method. Prior to vesicle formation, a lipid film of PPC 1:1:1 was prepared. On a pre-cleaned coverslip chamber, the vesicle fusion was assisted using 10 mM $Ca^{2+}$. The chamber preparation and cleaning procedure are already discussed elsewhere[9]. The prepared bilayer was incubated with 0.6 nM of RAβ oligomers for 30 minutes, washed with buffer, and finally imaged in the TIRF for membrane attachment. For measuring the attachment, movies were not recorded till complete bleaching, rather images of different ROIs were captured. For detecting the oligomer stoichiometry, movies of ROIs were recorded until the fluorescent spots/oligomers underwent complete bleaching.

Analysis of the movies was done using Fiji (Image-J-win64, freely available image analysis program). For detecting the oligomers, a particular threshold was provided and spots with intensity less than that were neglected. For a single spot, the diameter was restricted to 3×3 pixels (pixel size, 156 nm. To determine the stoichiometry, each spot was selected using the selection tool in Fiji and an intensity vs frames (i.e., time) plot was generated. The number of steps in which the fluorescence bleached in each spot was manually counted. This number gave the oligomer stoichiometry. A minimum of 300 spots were detected to assign the stoichiometry distribution in each case.

**SI 14: Titration of short-lifetime component in RN46A cells with different LVFFA concentration and cellular uptake**

To establish the importance of ApoE and RAβ oligomers, LVFFA titration was performed in RN46A cells. Briefly, LVFFA (DMSO stock) of different concentration were incubated in media containing cells for 1 hour. The amount of DMSO was maintained to be less than 1%. Post incubation, excess LVFFA was washed off using TB (thrice) and cells were incubated with RAβ oligomers for 3 hours. FLIM imaging was performed on these cells. As a sham control, cells were incubated with DMSO (1%) followed by incubation with RAβ oligomers.

The average lifetime distribution from the different samples (with different LVFFA concentration) were fitted using three Gaussians. The area under the Gaussian with the short-lifetime population was accounted to calculate the short-lifetime percentage upon LVFFA titration. Here we take a note that the short-lifetime seems slightly longer (around 1 ns) in these cells, yet much smaller than the lifetime of freshly prepared oligomers or free dye in RN46A cells. The lifetime distribution in presence of different LVFFA concentration is provided in figure S13A.

As a precaution, we measured whether the total cellular uptake of RAβ oligomers remains constant with LVFFA titration. Thus, RN46A cells treated with different LVFFA concentrations were imaged using confocal (Zeiss CLSM 880) and the average intensity from the cells was measured using ImageJ. Results showed that the uptake did not change with different LVFFA treatments (figure S13B).

**SI 15: Standardization of Aβ induced ROS measurements in RN46A cells**

To estimate the intracellular toxicity of the Aβ oligomers, reactive oxygen species (ROS) was measured in cells. High concentration stock of unlabelled Aβ peptide (1 mM, pH 11) was used to estimate the cell ROS at its different concentrations. To identify the concentration at which Aβ shows the maximum cell stress, cell ROS was measured for three different concentrations. 500 pM, 3 µM, and 5 µM concentrations of Aβ were used to check the cell ROS activity in RN46A cells. For the measurement, Aβ was incubated to the cells in media for 30 mins. Post incubation, the excess Aβ was washed off thrice using TB and cell ROX deep red (Excitation: 633 nm, Emission: 641 nm – 700 nm) was incubated for another 30 mins. As a sham control, Aβ was replaced with buffer and the incubation times were kept constant. The cell ROS was quantified by monitoring the average fluorescence intensity from z-projections of the cells.

We found with increasing concentration of Aβ, the cell ROS increases (figure S14). However, the cell morphology got severely compromised at 5 µM Aβ. Thus we carried out all further toxicity measurements of Aβ and dose dependence with LVFFA at 3 µM concentration.

**SI 16: Less toxic Aβ mutants shows negligible short-lifetime component in RN46A cells**

The F19-L34 residue contact in Aβ peptide is crucial for its toxicity[1]. It has been shown previously that minimal perturbation in this contact can drastically reduce the toxicity associated with Aβ[11]. One such Aβ variant is AβF19Cha. The phenylalanine in the 19$^{th}$

position is modified to cyclohexylalanine (an artificial amino acid without aromatic phenyl ring). This was expected to petFurb the F19-L34 contact as cyclohexyl group would prefer a non-planar conformation unlike phenyl group, thus perturbing the contact length. It was found that the toxicity of the peptide is significantly lower than WT Aβ[11]. Thus we checked whether in RN46A cells, a less toxic variant of Aβ shows a similar ApoE induced lifetime modification. Figure S15 shows the lifetime image and average lifetime distribution. The development of short-lifetime component (with similar experimental protocol as WT RAβ oligomers, 350-500 nM incubation for 3 hours) was negligible (6.7 ± 2.1%).

**SI 17: Among several proteins, ApoE 4 can introduce short-lifetime in RAβ oligomers *in-vitro* despite stable complex formation**

To check whether purified ApoE can introduce short-lifetime component in RAβ oligomers (100 nM), 1 µM DMPC lipidated ApoE[12] was incubated in TB at 37 °C. Lifetime measurement of the sample showed a short-lifetime component of 3.4% (data in main text). This was negligible as compared to the intracellular ApoE, however both lipidation state and source of ApoE are pivotal to determine the function of ApoE.

Fluorescence correlation spectroscopy (FCS) analysis (figure S16B) showed that the diffusion time ($\tau_D$) did not increase proportionately to the size of a stable complex between ApoE (34-36 Kda) and RAβ oligomers. MEM distribution showed only a slight increase in the diffusion time. This indicated a dynamic/labile interaction between ApoE and RAβ oligomers. The data also corroborated with the increase in oligomer stoichiometry as obtained from the smPB measurements (figure 2E 1, main text).

Both lifetime and FCS measurements of RAβ oligomers were carried out after incubation with other proteins (relevant in AD). Ubiquitin (UF45W)[13], Cytochrome C[14,15], Myoglobin[16], and a control protein BSA were tested. A tryptophan mutant of Ubiquitin (UF45W) was used to check whether ubiquitin can also show lifetime quenching (as tryptophan quenches rhodamine B)[10] since it is known to interact with Aβ[13]. Also it has been already established that the tryptophan point mutation do not hamper the physical and chemical properties of ubiquitin[17]. MEM distribution of the lifetime measurements showed no short-lifetime component with any other protein incubation, however in FCS measurements Ubiquitin showed binding (figure S16B). This again established that the short-lifetime in RAβ oligomers is ApoE specific.

### SI 18: Generation of neural stem cells (NSCs) and experimental details

#### A. Generation of NSCs

Patient-derived HiPSCs (Cell line No.: 8904, Late-onset AD with *APOE 3/4*), were obtained from the ADBS stem cell repository (https://www.ncbs.res.in/adbs/bio-repository). HiPSCs were cultured as feeder-free colonies on a Matrigel matrix (354277; Corning) in Essential-8 Media (A15169; Gibco). Media changes were given every 24-48 hours. Cells were passaged using StemPro™ Accutase™ (A1110501; Gibco), and re-plated at 1:3 dilutions. NSC generation was performed using our previously established Embryoid body (EB) based protocol with slight modifications[18]. Briefly, sub-confluent healthy HiPSC cultures were transferred from complete E8 media to a 1:1 combined media, consisting of E8 and EB media (KO-DMEM (10829018; Gibco), 20% knock-out serum replacement (10828-028; Gibco), 1x MEM-NEAA (11140050; Gibco), 1xGlutaMax (35050061; Gibco) and 1xPen-Strep (15140122; Gibco), supplemented with 0.1mM beta-mercaptoethanol (21985023; Gibco)) for 4-8 hours. The cells were lifted using StemPro™ Accutase™ and grown in low attachment dishes to form self-aggregating EBs in the above media supplemented with RevitaCell™ Supplement (A2644501; Gibco) for 24 hours. The following day, EBs were transferred to fresh low attachment dishes in complete EB media and cultured in suspension until day 6 with alternate day media change.

The growing EBs were transferred to neural induction media (DMEM/F12 (11320033; Gibco), supplemented with 20ng/mL bFGF (PHG6015; Gibco), 1x N2-supplement (17502-001; Gibco), 1x MEM-NEAA, 1x GlutaMax, 1xPen Strep and 2 µg/ml Heparin (H3149; Sigma)) on day 7 and cultured till day 14 with alternate day media change. Neuro-ectoderm-induced EBs were plated on Matrigel-coated adherent dishes for primary rosette formation in NIM on day 14. Rosettes were propagated manually for at least three rounds to enrich for neural epithelial cells and then plated on Laminin (23017015; Gibco) coated dishes in Neural expansion media (NEM) comprising of DMEM/F12, supplemented with 1x N2 supplement, 1x B27 supplement without RA (12587-010; Gibco), 20ng/ml bFGF, 1x MEM-NEAA, 1x GlutaMax, 1xPen Strep and 2 μg/ml Heparin. Further propagation of NSCs were done on Laminin-coated dishes in NEM.

To verify that our cultures were indeed a neural stem cell population (phase image in figure S17), we performed standard ICC using bona fide NSC marker antibodies, SOX2 (ab97959; Abcam), and NESTIN (ab22035, Abcam). Briefly, cells were grown on glass coverslips and fixed with 4% paraformaldehyde for 10 minutes, washed with 1xPBS, permeabilized with 0.1% TritonX-100 (CAS # 9002-93-1; Sigma-Aldrich), and blocked using 3% bovine serum albumin (A9418; Sigma-Aldrich). Primary antibodies were incubated on fixed cells overnight at 4ºC, washed thrice with 1xPBST, and then incubated with Alexa-Fluor secondary antibodies having excitation at 488nm and 594nm (Cat. Nos. - A11034 and A11005; Invitrogen) for 1 hour at room temperature. After three washes, the cells were incubated with DAPI (R37606; Invitrogen) for nuclear staining and imaged using Olympus IX-71 Inverted Epifluorescence Microscope, at 20x magnification.

**B. RAβ oligomers incubation experiments**

For fluorescence lifetime measurement experiments, the NSCs were seeded at a density of 0.04 x $10^6$ cells/$cm^2$ on Laminin-coated dishes. Media changes were given on alternate days after plating, until the fourth day. Cells were then incubated with 350nM R-$A\beta_{1-40}$ for 30 minutes at 37ºC. The media was removed, and cells were washed thrice with TB. Following this, to extract cell lysates, NSCs were gently scraped from the surface and centrifuged at 1200 rpm for 3 minutes. The cell pellet was re-suspended in 1% Triton-X 100 Lysis buffer (150mM Sodium Chloride (GRM853; HiMedia®) in 50mM Tris-HCl, pH 7.5 (15567027; Invitrogen™), containing Protease Inhibitor (A32953; Thermo Scientific™). The cell suspension was incubated on ice for 20 minutes with intermittent vortexing. The obtained lysate solution was centrifuged at 4000 rpm for 15 minutes at 4ºC. The supernatant was filtered through a 0.22μ filter and stored at 4ºC until further use.

Supplementary figure 1

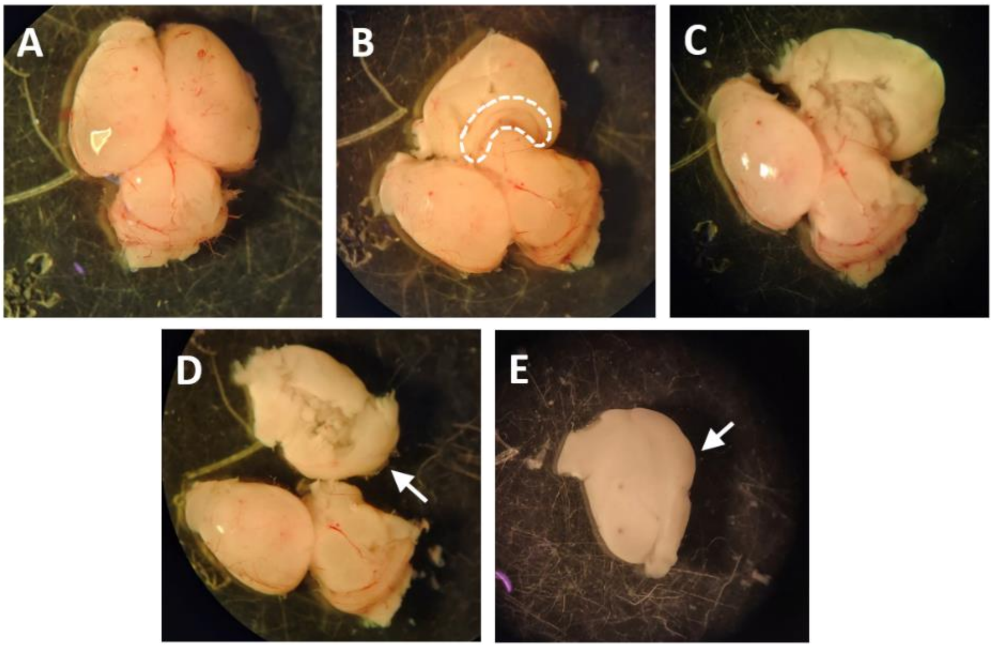


**Figure S1:** Dissection of cortex from the brain hemispheres of Sprague Dawley pups. (A) The entire brain of a P2 pup (with meninges), (B) Separation of the brain hemispheres results in exposure of the cortex and the hippocampus. The marked region is the hippocampus in the particular hemisphere. (C) Removal of the hippocampus, (D) Separation of the entire cortex from the brain, and (E) Demeningized cortex.

Supplementary figure 2

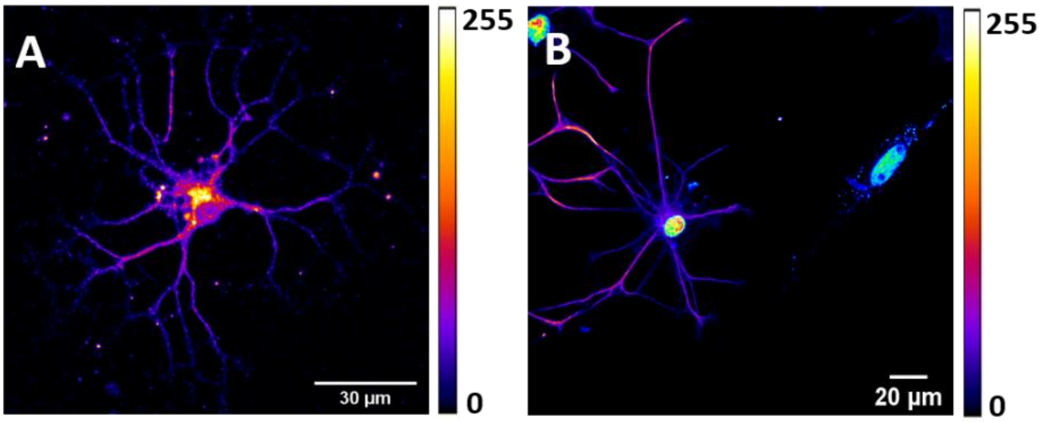


**Figure S2:** Isolation and detection of astrocytes from the cortical culture. (A) ICC of a single astrocyte using antibody targeted against GFAP. Intensity is false colour-coded for the Atto 488 label in the secondary antibody, (B) In a cortical culture, selectively the astrocytes are labelled over cells. Clearly an adjacent fibroblast is not labelled

Supplementary figure 3

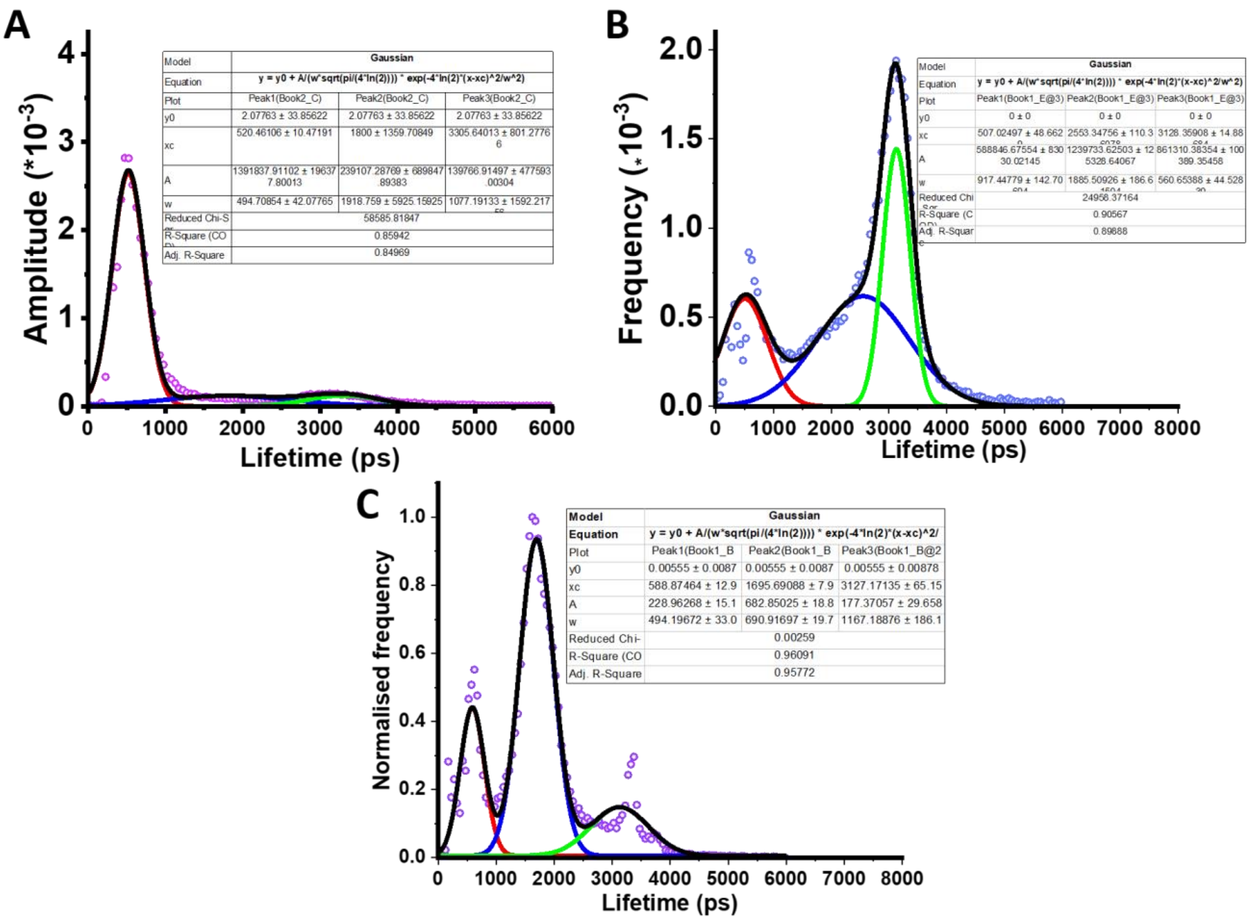


**Figure S3:** Average lifetime distribution of RAβ oligomers from cells. (A) Distribution in astrocytes, (B) distribution in cortical neurons, and (C) distribution in RN46A cells. The red, blue, and green lines are the Gaussian fits for the first, second, and third components respectively. Black is the cumulative fit of all the Gaussians.

Supplementary figure 4

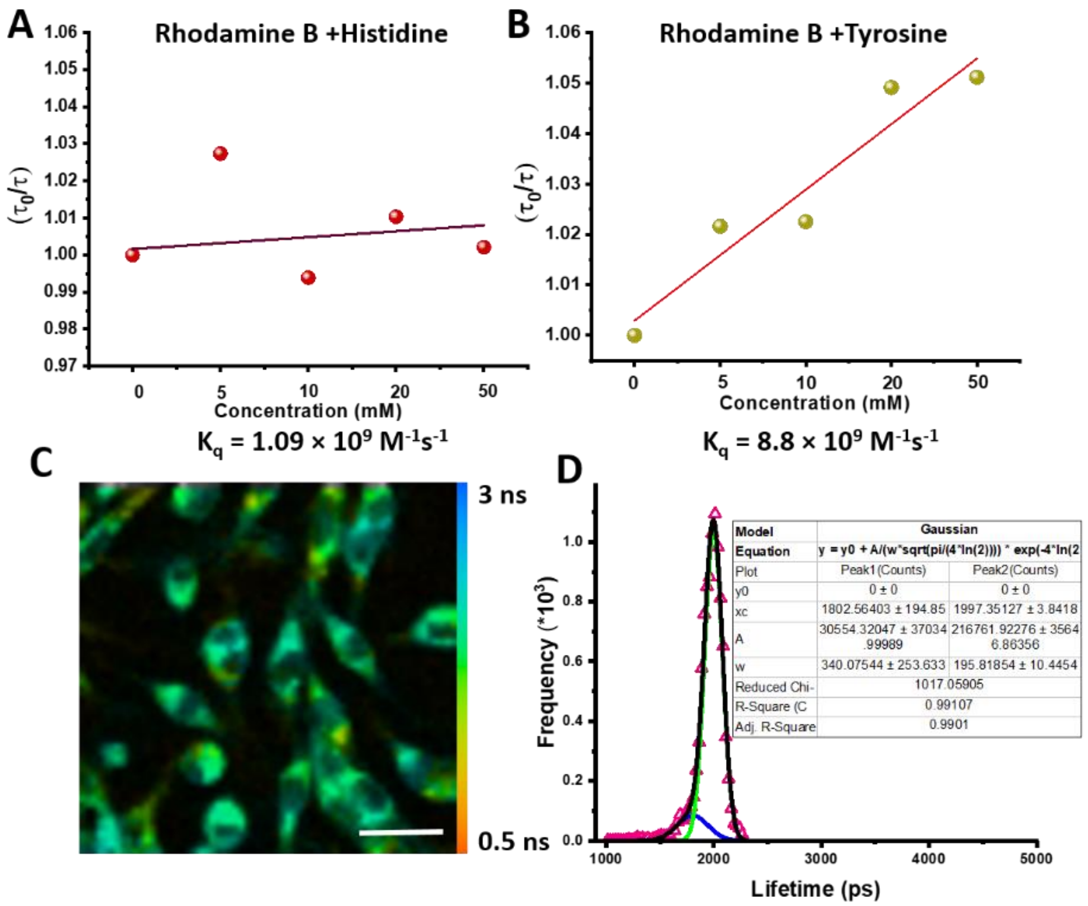


**Figure S4:** Lifetime quenching of 100 nM rhodamine B dye by histidine and arginine. (A) quenching by histidine, (B) quenching by l-tyrosine (pH-9). (C) Lifetime image of rhodamine B dye in RN46A cells, and (D) Average lifetime distribution of rhodamine B in RN46A cells. The blue, and green lines are the Gaussian fits for the two distinct lifetime components. Black is the cumulative fit of the two Gaussians. The fit parameters are given in the box. (scale bar in fig 4C is 30 µm).

Supplementary figure 5

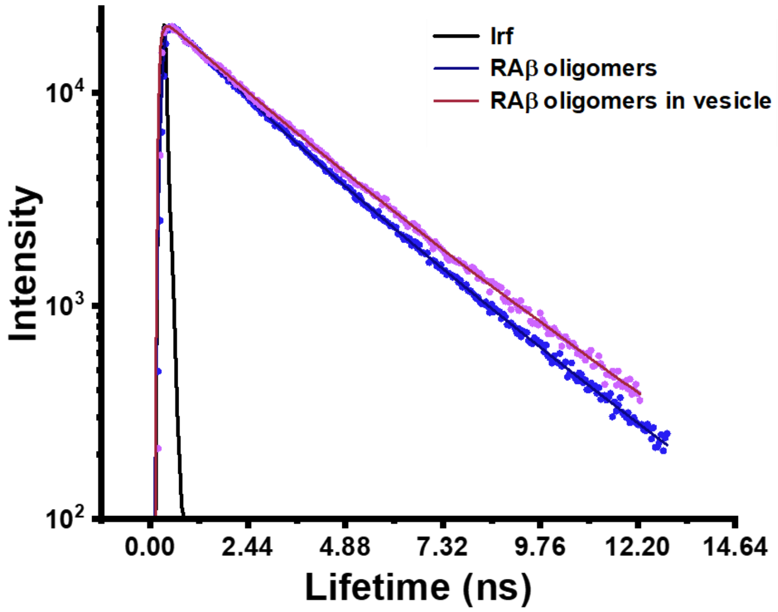


**Figure S5:** Lifetime decay of RAβ oligomers in solution and PPC 111 vesicles. Table 9 shows the lifetime components and the average lifetime decay values.

Supplementary figure 6

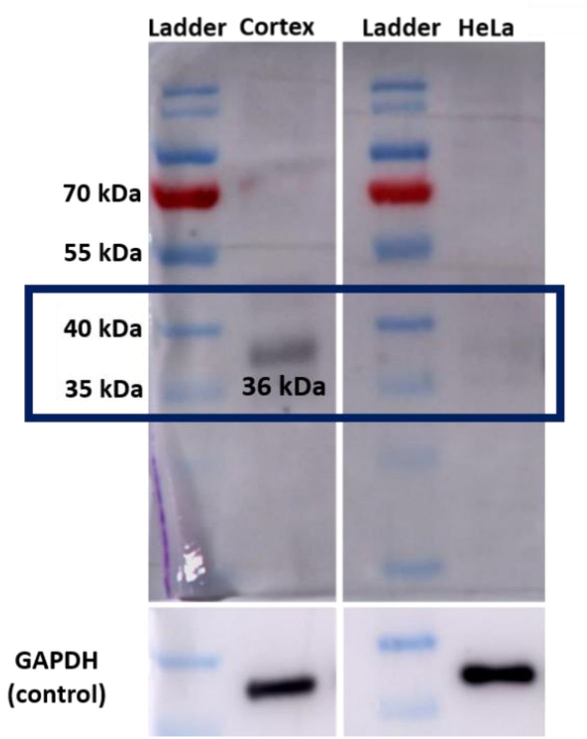


**Figure S6:** Western blotting of EPR19392 anti-ApoE antibody with full brain cortex and HeLa cells. GAPDH was used as an internal control. A 36 kDa band shows the presence of ApoE in the cortex extract and was not detected in the HeLa cells.

Supplementary figure 7

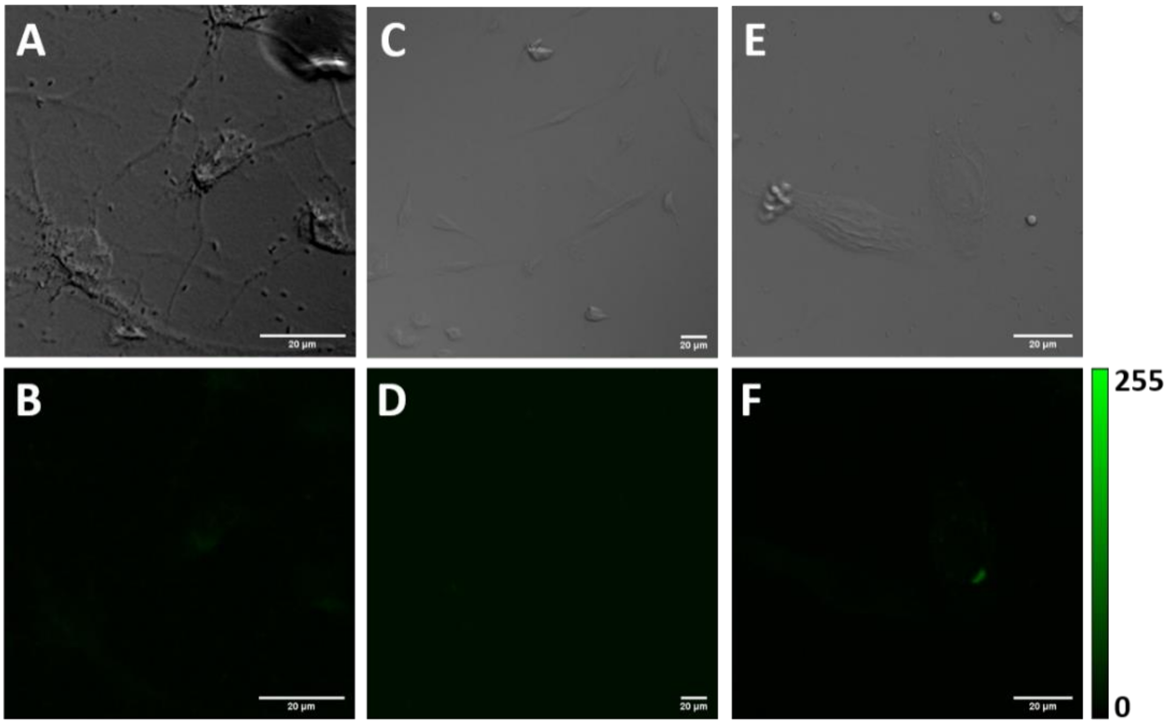


**Figure S7:** Control ICCs for ApoE in cells (without the incubation of primary antibodies). A relatively background free ICC indicates negligible error in estimating the intracellular ApoE across different cells. Transmission and Intensity of (A, B) astrocytes, (C, D) RN46A cells, and (E, F) HeLa cells.

Supplementary figure 8

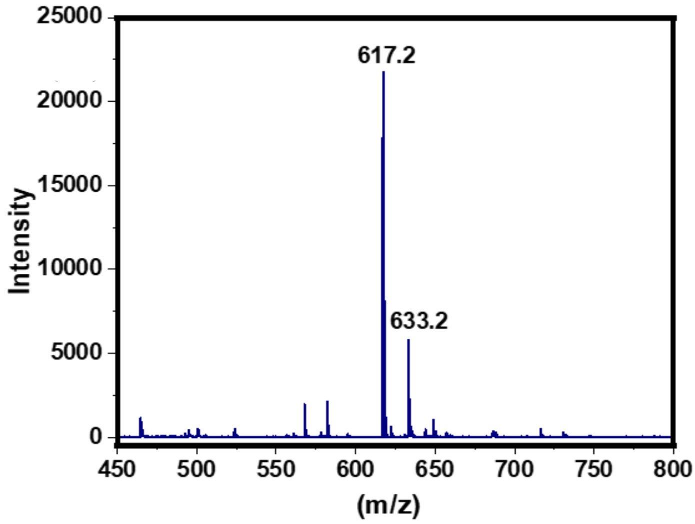


**Figure S8:** MALDI mass spectrum of the LVFFA. The $Na^+$ and $K^+$ adduct peaks are visible in the spectrum. For experiment, the LVFFA peptide powder was dissolved in DMSO and a 10 mM stock solution was prepared. For lifetime experiments, the peptide (of required concentration) was incubated in the cells (in TB) for 1 hours. RAβ oligomers (350-500 nm) was directly added to the buffer for incubation without any intermittent washing step. After the desired incubation time, excess RAβ oligomers and LVFFA washed off and cells were imaged.

Supplementary figure 9

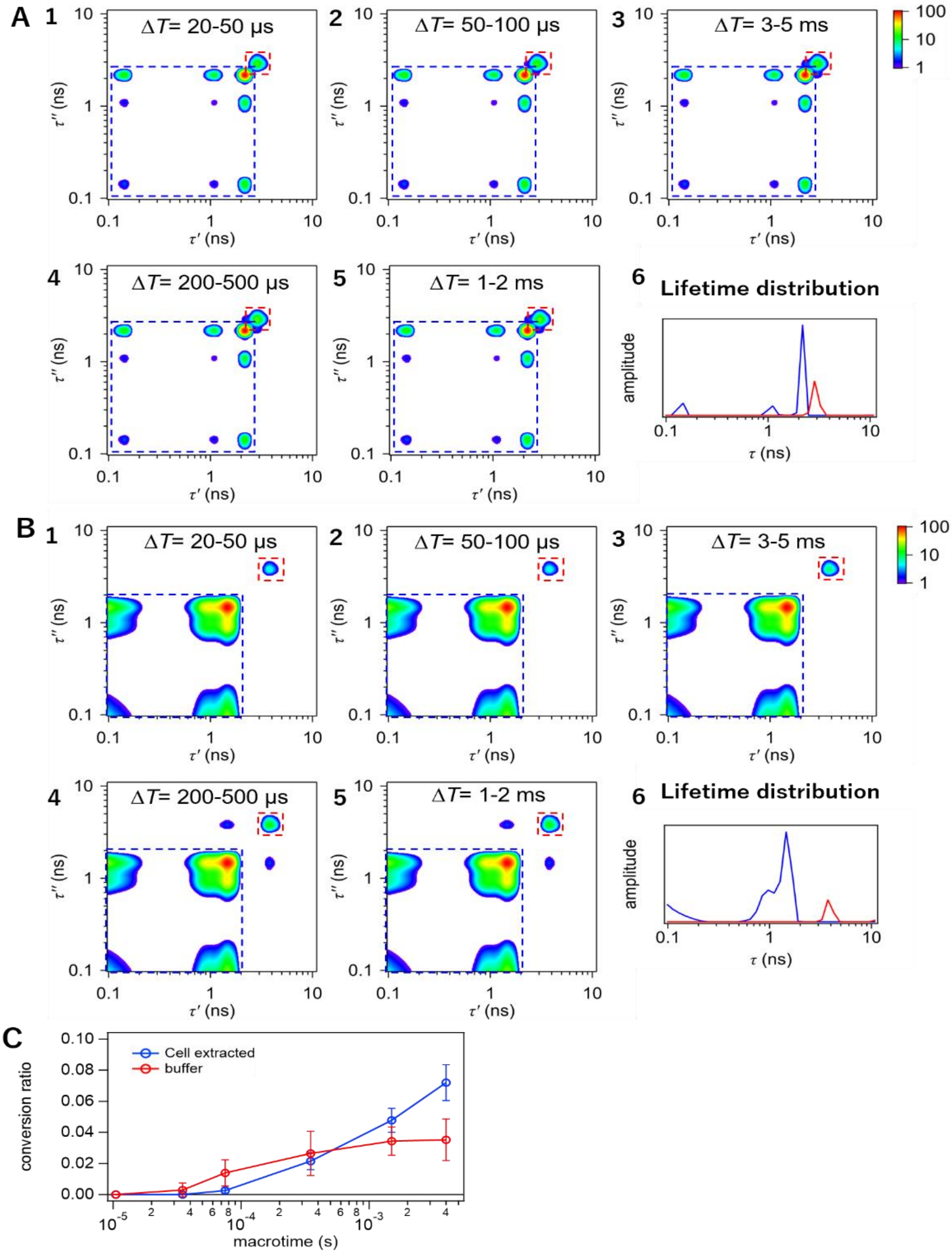

**Figure S9:** 2D FLCS characterization of the RAβ oligomers prepared freshly in TB and extracted from cortical extracts. **A (1-5)** rest 2D emission decay maps of freshly prepared RAβ oligomers at different macrotime intervals and, **(6)** the lifetime distribution obtained from global analysis of 2D FLCS curves. Blue is the short lifetime and red is the longer lifetime. **B (1-5)** 2D emission decay maps of RAβ oligomers extracted from cortical culture at different macrotime intervals and, **(6)** the lifetime distribution obtained from global analysis of 2D FLCS curves. **C** Interconversion dynamics of the two distinct species of RAβ oligomers in TB and extracted from cortical culture. As defined previously and can be seen from the data, interconversion dynamics between the different species is negligible. Values indicate mean ± standard deviation from 2D FLCS analysis.

Supplementary figure 10

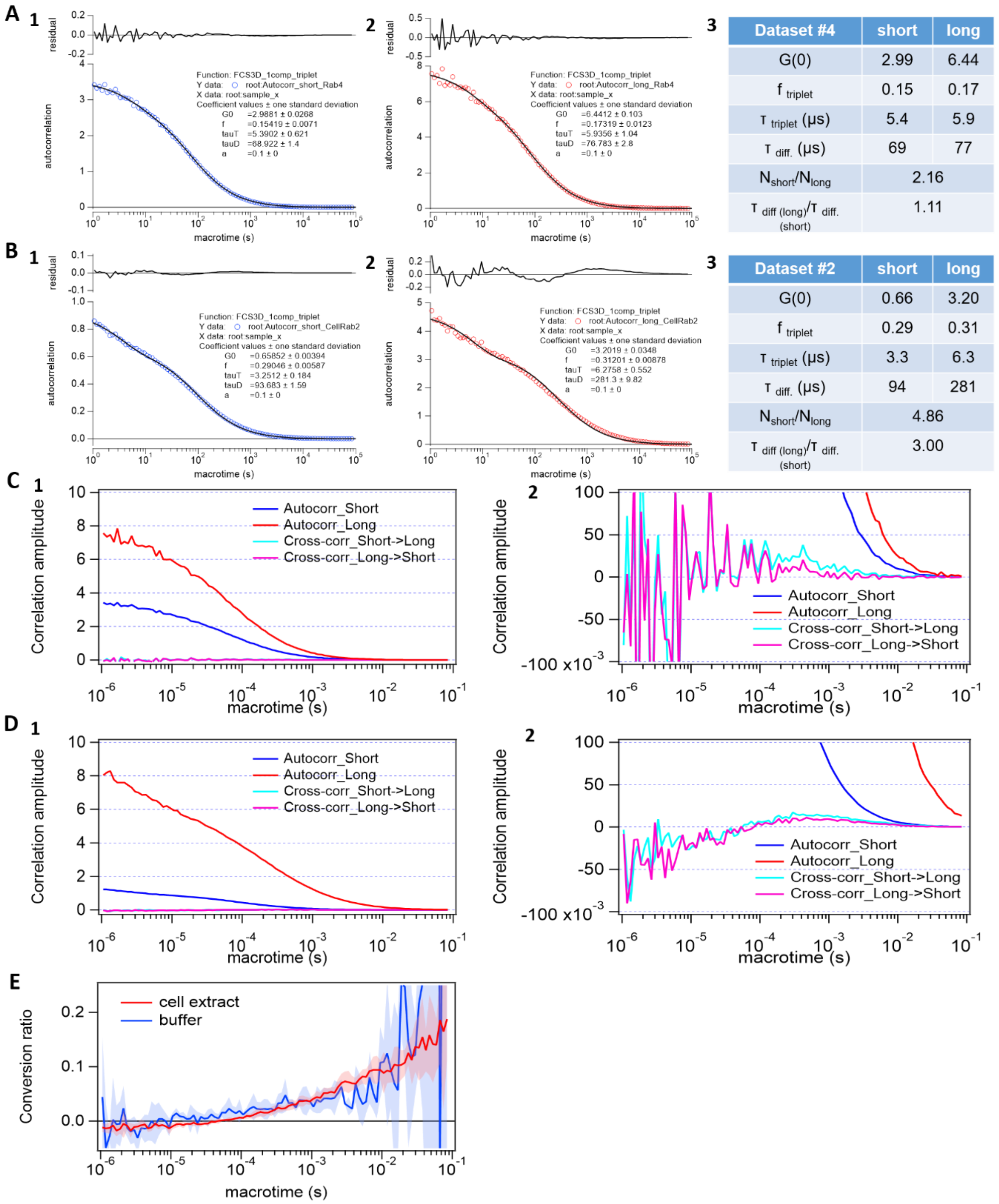


| Dataset #4 | short | long |
|---|---|---|
| G(0) | 2.99 | 6.44 |
| f triplet | 0.15 | 0.17 |
| τ triplet (μs) | 5.4 | 5.9 |
| τ diff. (μs) | 69 | 77 |
| $N_{short}/N_{long}$ | 2.16 | |
| $\tau_{diff\ (long)}/\tau_{diff.\ (short)}$ | 1.11 | |

| Dataset #2 | short | long |
|---|---|---|
| G(0) | 0.66 | 3.20 |
| f triplet | 0.29 | 0.31 |
| τ triplet (μs) | 3.3 | 6.3 |
| τ diff. (μs) | 94 | 281 |
| $N_{short}/N_{long}$ | 4.86 | |
| $\tau_{diff\ (long)}/\tau_{diff.\ (short)}$ | 3.00 | |

**Figure S10:** Global species specific auto-correlation of the RAβ oligomers in buffer **A**, and extracted from cortical culture **B**. Blue and red traces represent the short and long lifetimes respectively. Parameters for each are given in the adjacent tables. Filtered FCS analysis of state specific auto and cross-correlation of freshly prepared RAβ oligomers (**C1**, and **2** - vertically expanded in the y-axis), and RAβ oligomers extracted from cortical culture (**D1**, and **2** - vertically expanded in the y-axis). **E** Conversion ratio obtained from the filtered FCS analysis. Solid curves and shaded regions indicate mean ± standard deviation from filtered FCS analysis.

Supplementary figure 11

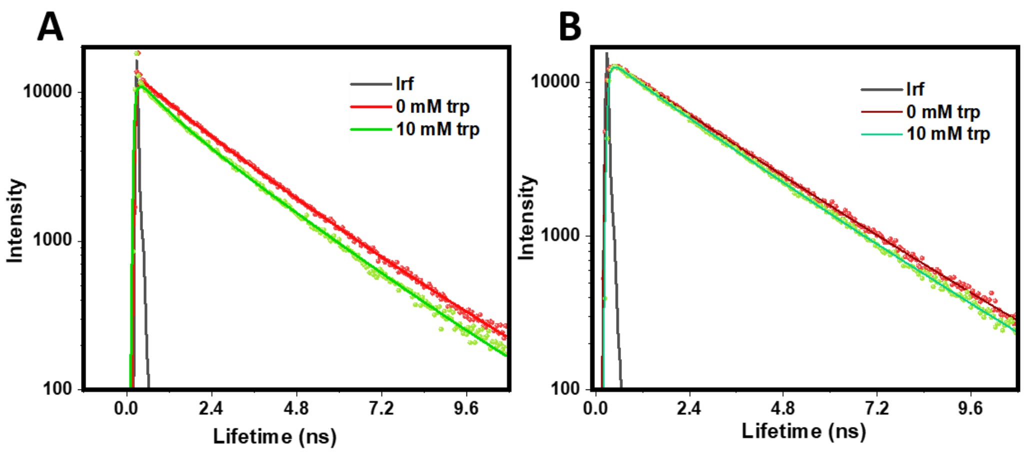


**Figure S11: A** TCSPC decays of RAβ oligomers with tryptophan (lifetime quenching measurement) (A) extracted from RN46A cells, and (B) freshly prepared (average lifetime). Table S15 shows the fit parameters and lifetimes of cell extracted RAβ oligomers with different tryptophan concentration, and S16 shows the same for freshly prepared RAβ oligomers.

Supplementary figure 12

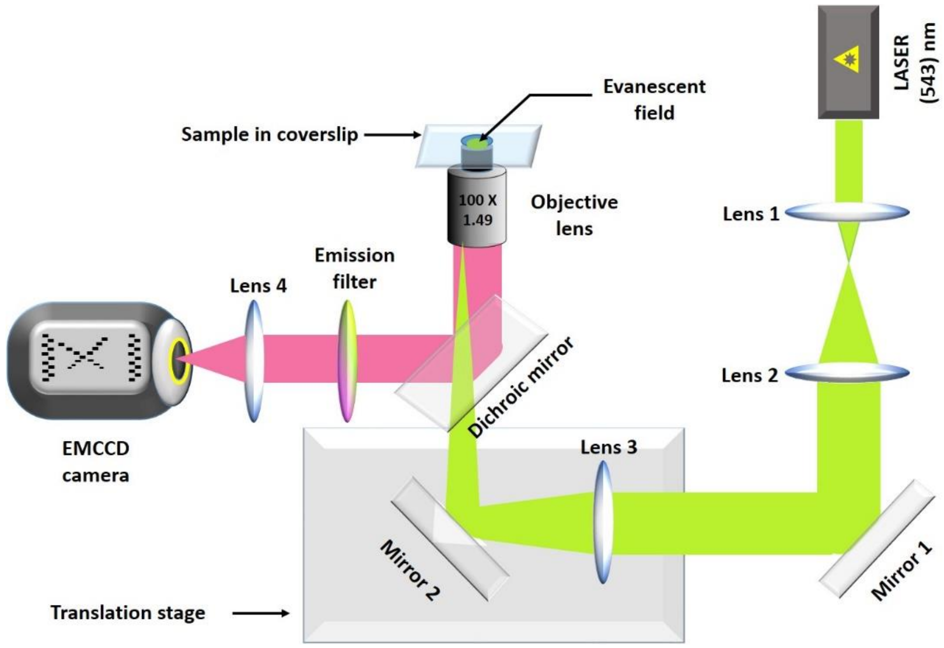


**Figure S12:** A diagrammatic representation of the home-built TIRF set up. Lenses 1 and 2 form a telescope that increases the diameter of the excitation laser beam (green). Lens 3 focuses the beam at the back focal plane of the objective lens. The translation stage is used to adjust the penetration depth, as needed. The fluorescence (red) is separated from the excitation beam using a dichroic mirror. The fluorescence is passed through an emission filter before it is focused on to an EMCCD camera using the tube lens (lens 4).

Supplementary figure 13

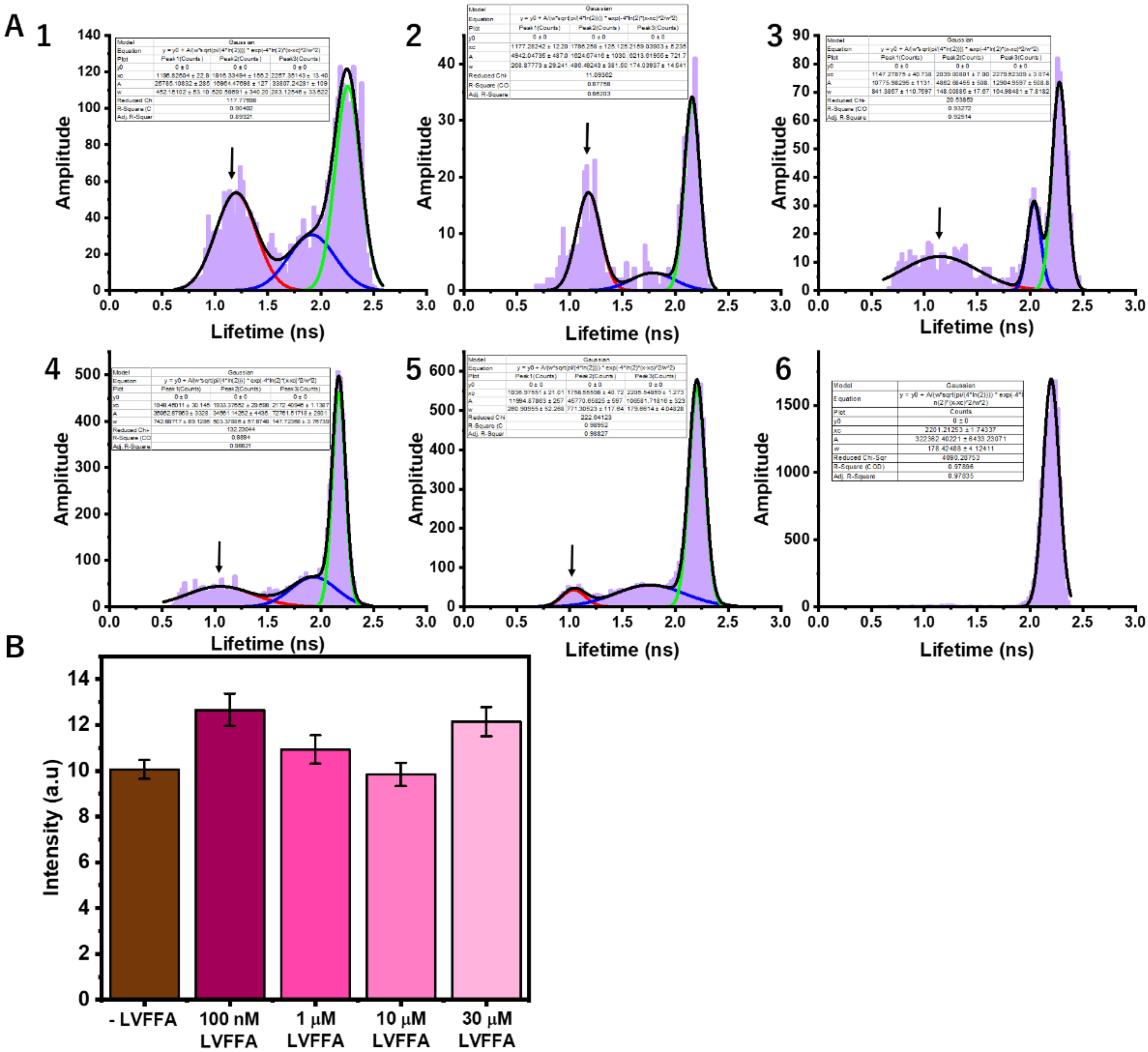


**Figure S13: A** Lifetime distribution of RAβ oligomers in RN46A cells in presence of different LVFFA concentration, (1) sham control, (2) 0.1 µM, (3) 0.5 µM, (4) 1 µM, (5) 10 µM, and (6) 30 µM respectively. The short-lifetime component is marked using an arrow, and **B** total intracellular uptake of RAβ oligomers in presence of different LVFFA concentration.

Supplementary figure 14

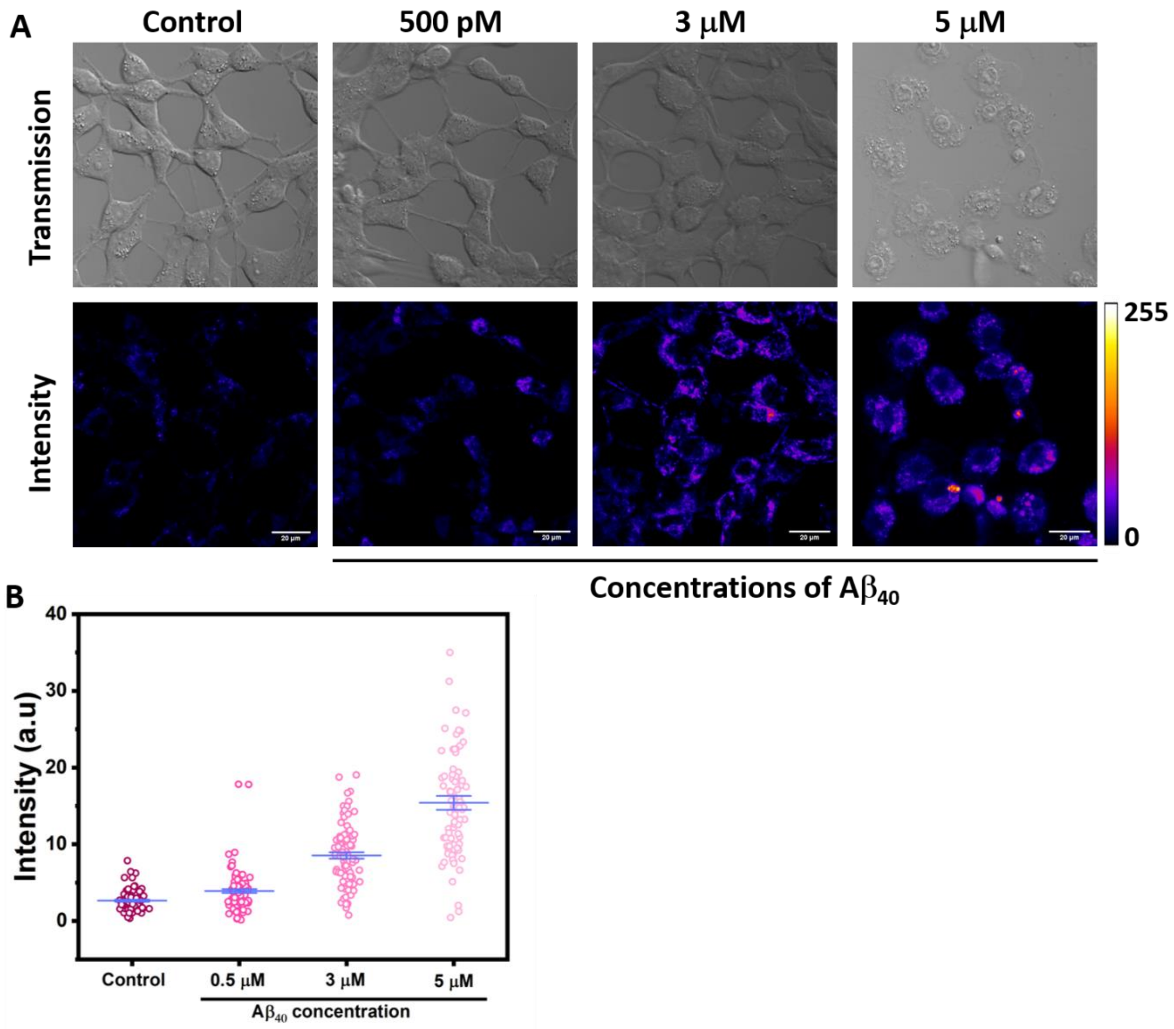


**Figure S14: A** ROS Intensity and transmission images of RN46A cells with sham control, (0.5 μM Aβ, 3 μM Aβ, and 5 μM Aβ. **B** Quantification of the ROS intensity. Data plotted is mean and standard error from three independent measurements (three dishes each measurement).

Supplementary figure 15

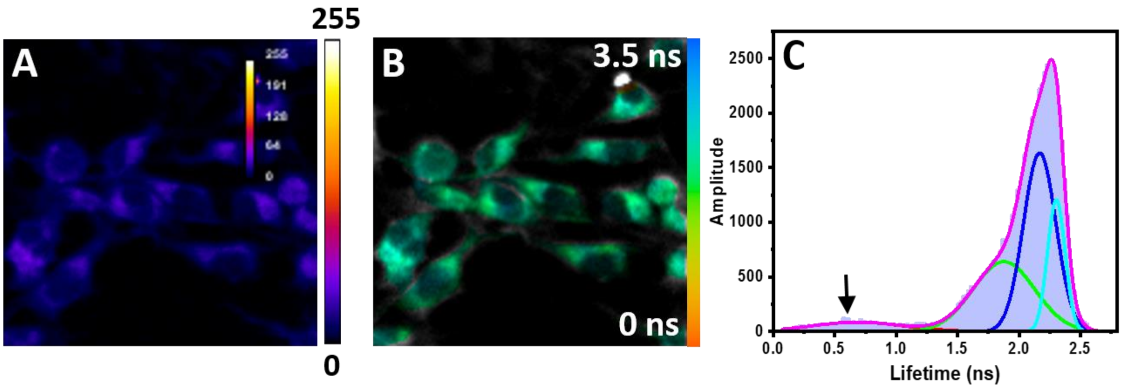


**Figure S15:** FLIM imaging of RaβCha oligomers in RN46A cells **A** Intensity image, **B** Lifetime image, and **C** Average lifetime distribution of RaβCha oligomers. The short-lifetime is marked by an arrow. Area quantification yielded a negligible short-lifetime component.

Supplementary figure 16

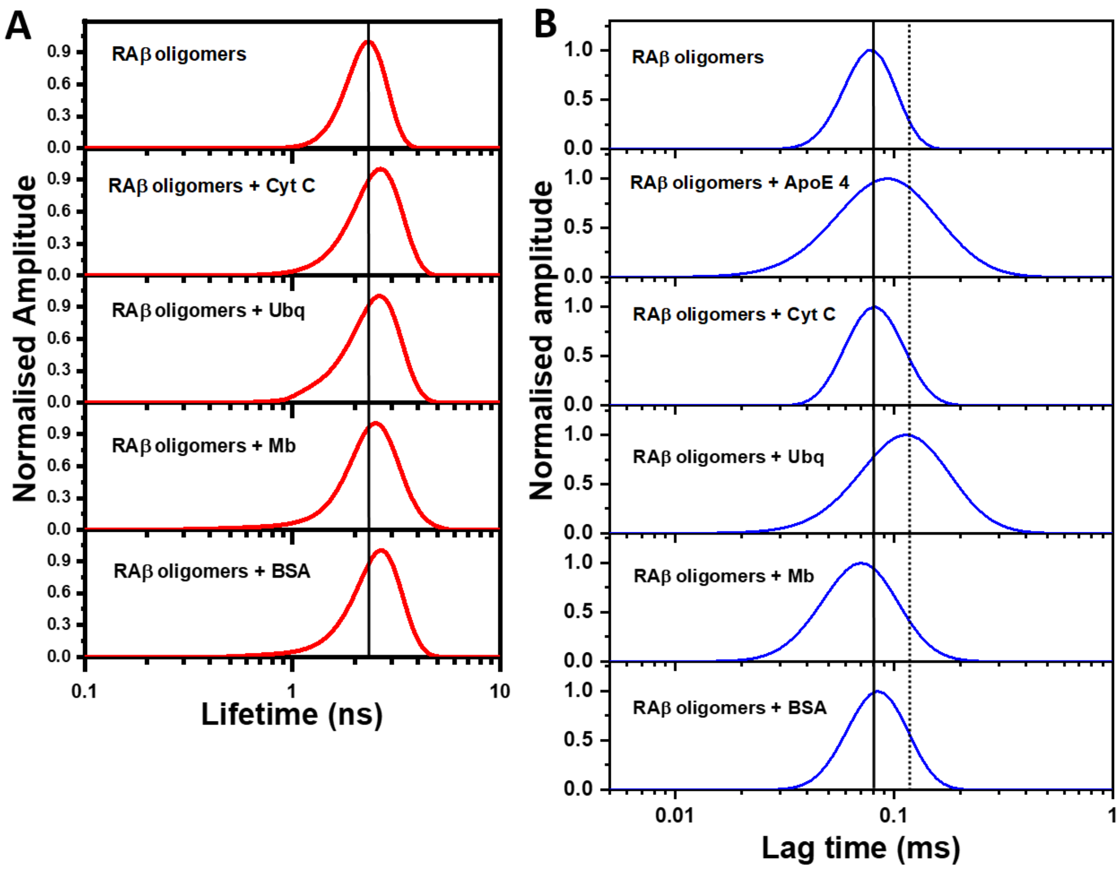


**Figure S16: A** MEM lifetime distribution of RAβ oligomers freshly prepared, incubated with 5 µM Cytochrome C, 10 µM Ubiquitin (UF45W), 5 µM Myoglobin, and 5 µM BSA. The horizontal line is a guide to the eye of the lifetime maximas. **B** MEM FCS distribution of RAβ oligomers and proteins as mentioned previously of the same concentration. The solid line represents the diffusion time ($\tau_D$) of freshly prepared RAβ oligomers (80 µs). Diffusion time in presence of UF45W is more than 100 µs (longer than the ApoE 4 incubated sample).

Supplementary figure 17

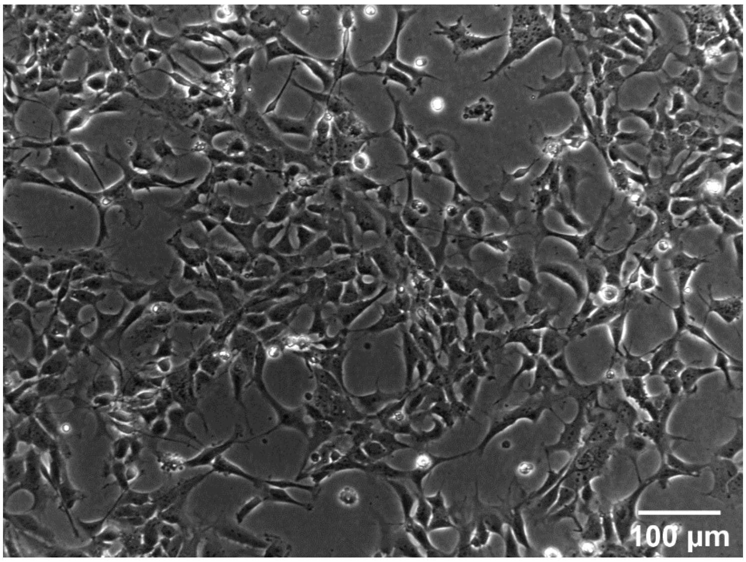


**Figure S17:** Phase image of NSC (cell line number 8904, with ApoE 3/4)

Supplementary tables

| Components | Volume (ml) |
|---|---|
| 10×HBSS | 50 |
| 1M Hepes (pH 7.3) | 6.25 |
| 22.5% Glucose | 70 |
| Pen-Strep(100 ) | 10 |

**Table S1:** Composition of HHGN buffer.

| Components | Amount |
|---|---|
| NaCl | 800 mg |
| KCl | 400 mg |
| $KH_2PO_4$ | 60 mg |
| $Na_2HPO_4$ | 48 mg |
| $NaHCO_3$ | 350 mg |
| Glucose | 1000 mg (not to be added for HHGN) |

**Table S2:** composition of HBSS buffer.

| Components | Concentration (mM) | Weight (gm) |
|---|---|---|
| NaCl | 146 | 0.4260 |
| KCl | 5.4 | 0.0201 |
| $CaCl_2.2H_20$ | 1.8 | 0.0132 |
| $MgSO_4$ | 0.8 | 0.0099 |
| $KH_2PO_4$ | 0.4 | 0.0027 |
| $Na_2HPO_4$ | 0.3 | 0.0027 |
| d-Glucose | 5 | 0.0450 |

| | | |
|---|---|---|
| Na Hepes | 20 | 0.2603 |

**Table S3:** Composition of Thomson's buffer (T.B).

| Components | Volume (ml) |
|---|---|
| DMEM high glucose | 50 |
| FBS | 10 |
| Neurobasal | 40 |
| B-27 supplement | 2 |
| 10x pen-strep | 1 |

**Table S4:** Composition of primary culture media.

| Cell type | Population 1 mean lifetime (ps) | Contribution | Population 2 mean lifetime (ps) | contribution | Population 3 mean lifetime (ps) | contribution |
|---|---|---|---|---|---|---|
| **Astrocytes** | 520.46 | 0.78 | 1800.0 | 0.135 | 3305.64 | 0.078 |
| **Cortical neurons** | 507.02 | 0.21 | 2553.34 | 0.46 | 3128.35 | 0.32 |
| **RN46A** | 588.87 | 0.21 | 1695.69 | 0.62 | 3127.17 | 0.16 |

**Table S5:** Lifetime and population from independent fits of FLIM data

| Model | Gaussian | | |
|---|---|---|---|
| Equation | y = y0 + A/(w*sqrt(pi/(4*ln(2)))) * exp(-4*ln(2)*(x-xc)^2/w^2) | | |
| Plot | Peak1(B) | Peak2(B) | Peak3(B) |
| y0 | 0 ± 0 | 0 ± 0 | 0 ± 0 |
| xc | 549.2884 ± 3.153 | 1610 ± 193.96183 | 3303.53468 ± 174.4 |
| A | 324.83652 ± 13.1 | 68.02771 ± 29.6332 | 38.56916 ± 16.9828 |
| w | 430.93762 ± 11.6 | 1686.42587 ± 861.3 | 1063.26535 ± 347.6 |
| Reduced Chi- | 4.18486E-4 | | |
| R-Square (CO | 0.98173 | | |
| Adj. R-Square | 0.98061 | | |

**Table S6:** Global fit parameters for FLIM of RAβ oligomers in astrocytes.

| Model | Gaussian | | |
|---|---|---|---|
| Equation | y = y0 + A/(w*sqrt(pi/(4*ln(2)))) * exp(-4*ln(2)*(x-xc)^2/w^2) | | |
| Plot | Peak1(Book1_C) | Peak2(Book1_C) | Peak3(Book1_C) |
| y0 | 0 ± 0 | 0 ± 0 | 0 ± 0 |
| xc | 604.50124 ± 18.255 | 2369 ± 63.2778 | 3123.97705 ± 6.544 |
| A | 205.10883 ± 15.061 | 565.90241 ± 35.22699 | 515.95199 ± 26.306 |
| w | 667.33867 ± 46.958 | 1913.93058 ± 104.947 | 606.42014 ± 20.209 |
| Reduced Chi-Sq | 0.00135 | | |
| R-Square (COD) | 0.97985 | | |
| Adj. R-Square | 0.97838 | | |

**Table S7:** Global fit parameters for FLIM of RAβ oligomers in cortical neurons.

| Model | Gaussian | | |
|---|---|---|---|
| Equation | y = y0 + A/(w*sqrt(pi/(4*ln(2)))) * exp(-4*ln(2)*(x-xc)^2/w^2) | | |
| Plot | Peak1(D) | Peak2(D) | Peak3(D) |
| y0 | 0 ± 0 | 0 ± 0 | 0 ± 0 |
| xc | 588.03858 ± 13.02 | 1694.64769 ± 7.95 | 3121.34772 ± 66.23 |
| A | 235.93883 ± 12.52 | 686.46548 ± 17.80 | 191.89373 ± 21.975 |
| w | 506.61546 ± 31.45 | 690.9286 ± 19.795 | 1226.46799 ± 172.4 |
| Reduced Chi-S | 0.00257 | | |
| R-Square (CO | 0.96079 | | |
| Adj. R-Square | 0.95796 | | |

**Table S8:** Global fit parameters for FLIM of RAβ oligomers in RN46A cells.

| Model | Gaussian | |
| --- | --- | --- |
| Equation | y = y0 + A/(w*sqrt(pi/(4*ln(2)))) * exp(-4*ln(2)*(x-xc)^2/ | |
| Plot | Peak1(Book1_E) | Peak2(Book1_E) |
| y0 | 0.00223 ± 0.00356 | 0.00223 ± 0.00356 |
| xc | 2400 ± 56.00846 | 2773.62241 ± 4.18208 |
| A | 291.33637 ± 23.55598 | 446.53297 ± 14.54811 |
| w | 1966.30089 ± 137.2779 | 536.25558 ± 12.44636 |
| Reduced Chi-Sqr | 6.62964E-4 | |
| R-Square (COD) | 0.98586 | |
| Adj. R-Square | 0.98522 | |

**Table S9:** Global fit parameters for FLIM of RAβ oligomers in HeLa cells are in table 9.

| Sample | $a_1$ | $t_1$ | $a_2$ | $t_2$ | $t_{avg}$ | $X^2$ |
| --- | --- | --- | --- | --- | --- | --- |
| **RAβ oligomers** | 0.35 | 1.59 | 0.65 | 2.87 | 2.44 | 1.05 |
| **RAβ oligomers in vesicle** | 0.37 | 1.96 | 0.63 | 3.19 | 2.86 | 1.15 |

**Table S10:** Lifetime parameters of RAβ oligomers in solution and vesicle

| Sample | $a_1$ | $t_1$ | $a_2$ | $t_2$ | $t_{avg}$ | $X^2$ |
| --- | --- | --- | --- | --- | --- | --- |
| **RAβ oligomers (RN46A extracted)** | 0.39 | 0.76 | 0.61 | 2.38 | 2.10 | 1.29 |
| **RAβ oligomers (RN46A extracted) + 20 mM EDTA** | 0.31 | 0.84 | 0.69 | 2.43 | 2.22 | 1.27 |

**Table S11:** Effect of 20 mM EDTA the cell extracted RAβ oligomers

| Sample | $a_1$ | $t_1$ | $a_2$ | $t_2$ | $t_{avg}$ | $X^2$ |
|---|---|---|---|---|---|---|
| **Extracted RAβ oligomers (pH 7.5)** | 0.38 | 0.71 | 0.62 | 2.38 | 2.12 | 1.17 |
| **Extracted RAβ oligomers (pH 4.5)** | 0.39 | 0.76 | 0.61 | 2.31 | 2.04 | 1.19 |

**Table S12:** Effect of different pH on the cell extracted RAβ oligomers

| Components | Amounts |
|---|---|
| Water | 2.7 ml |
| 30% Arylamide | 0.67 ml |
| 1.5 M Tris (pH-8.8) | 0.5 ml |
| 10% SDS | 40 µl |
| 10% APS | 40 µl |
| TEMED | 4 - 6 µl |

**Table S13:** Loading gel composition

| Components | Amounts |
|---|---|
| Water | 1.9 ml |
| 30% Arylamide | 1.7 ml |
| 1.5 M Tris (pH-8.8) | 1.3 ml |
| 10% SDS | 50 µl |
| 10% APS | 50 µl |
| TEMED | 4 - 6 µl |

**Table S14:** Resolving gel composition

| Cell extracted RAβ oligomers | $a_1$ | $t_1$ (ns) | $a_2$ | $t_2$ (ns) | t-avg |
|---|---|---|---|---|---|
| **0 mM trp** | 0.219 | 1.047 | 0.781 | 2.625 | 2.466 |
| **3 mM trp** | 0.215 | 1.045 | 0.785 | 2.619 | 2.464 |
| **5 mM trp** | 0.242 | 0.902 | 0.758 | 2.552 | 2.385 |
| **10 mM trp** | 0.283 | 0.534 | 0.717 | 2.435 | 2.284 |

**Table S15:** Lifetime quenching of RAβ oligomers (extracted from RN46A cells) with tryptophan

| Freshly prepared RAβ oligomers | t-avg |
|---|---|
| **0 mM trp** | 2.67 ns |
| **3 mM trp** | 2.64 ns |
| **5 mM trp** | 2.60 ns |
| **10 mM trp** | 2.53 ns |

**Table S16:** Lifetime quenching of RAβ oligomers (freshly prepared in T.B) with tryptophan

**SI 19: References**

(18) Mukherjee, O.; Acharya, S.; Rao, M. Making NSC and Neurons from Patient-

Derived Tissue Samples. *Methods Mol. Biol.* **2019**, *1919*, 9–24.
